# Endothelial deletion of ADGRF5 (GPR116) promotes fibro-inflammatory EndMT and impairs adaptive thermogenesis in brown adipose tissue

**DOI:** 10.1101/2025.10.31.683372

**Authors:** Rabih El-Merahbi, Vasiliki Karagiannakou, Ronja Kardinal, Lea Seep, Michelle Ynonne Jäcksein, Staffan Hildebrand, Mersiha Hasic, Eylül Korkmaz, Ankush Kumar Jha, Aspasia Thodou Krokidi, Ken Dyar, Felix Meissner, Stephan Grein, Jörg Heeren, Martin Klingenspor, Alexander Pfeifer, Jan Hasenauer, Dagmar Wachten, Stephan Herzig, Anastasia Georgiadi

## Abstract

**Objective:** Brown adipose tissue (BAT) dissipates energy via non-shivering thermogenesis, and it is a promising therapeutic target for metabolic disease. While most research focuses on thermogenic adipocytes, emerging data point to critical contributions from the surrounding stromal niche. Here, we investigated the role of adhesion G protein–coupled receptors (aGPCRs) in BAT function, focusing on Adgrf5 (Gpr116), a receptor enriched in endothelial cells.

**Methods:** We used single-nuclei RNA sequencing to map aGPCRs expression across mouse and human BAT. We then examined the consequences of Adgrf5(Gpr116) loss using global, brown adipocyte, and endothelial-specific knockout mouse models under acute and prolonged cold exposure.

**Results:** Inducible endothelial deletion of Adgrf5(Gpr116) impaired the maintenance of thermogenic capacity during prolonged—but not acute—cold exposure. This was not associated with defective angiogenesis, but rather with endothelial fibro-inflammatory reprogramming. Single-nuclei RNA sequencing analysis revealed endothelial-to-mesenchymal transition (EndMT) features, including induction of mesenchymal markers, collagens, and metalloproteinases, and loss of barrier genes. Adgrf5(Gpr116)-deficient endothelial cells also exhibited cytoskeletal remodeling and activation of stress fiber pathways, implicating Adgrf5(Gpr116) as a mechanosensory safeguard of endothelial identity.

**Conclusion:** Endothelial Adgrf5(Gpr116) preserves thermogenic competence in BAT by suppressing EndMT and maladaptive matrix remodeling. Our findings establish vascular mechanosensing as a critical determinant of thermogenic tissue homeostasis.

**Highlights:**

- Adhesion GPCRs are the second most abundant GPCR family in mouse and human brown fat
- Adhesion GPCRs are enriched in non-adipocyte cell types in brown fat and participate in cell–cell contact signaling
- Endothelial Adgrf5(Gpr116) is required for thermogenic adaptation during prolonged cold exposure
- Loss of Adgrf5(Gpr116) induces fibro-inflammatory reprogramming and endothelial-to-mesenchymal transition (EndMT)

## 1. Introduction

Brown adipose tissue (BAT) is a specialized thermogenic organ that dissipates energy as heat via non-shivering thermogenesis (NST). It has garnered increasing attention not only for its potential to reduce adiposity but also for its endocrine functions that improve systemic metabolism (1, 2). Functional BAT is present in adult humans and is activated by cold exposure or β-adrenergic stimulation. Importantly, BAT mass inversely correlates with the incidence of prediabetes, hyperlipidemia, and hypertension, suggesting a protective role in cardiometabolic health (3). Additionally, white adipose tissue (WAT) can acquire brown-like characteristics in response to environmental or hormonal cues, giving rise to beige adipocytes with thermogenic capacity (4). These findings underscore the therapeutic potential of enhancing BAT or beige fat activity to address obesity and related metabolic diseases.

Although functional BAT persists into adulthood, particularly in humans with lean phenotypes, its thermogenic activity appears to be reduced in aged individuals, people with obesity, and those with type 2 diabetes (5–10). Most research to date has focused on the thermogenic adipocytes themselves—their differentiation, mitochondrial uncoupling, and adrenergic responsiveness. However, brown adipocytes reside in a structurally and functionally integrated microenvironment, embedded in a dense extracellular matrix (ECM) that includes collagens, proteoglycans, adhesion proteins, and enzymes such as matrix metalloproteinases (MMPs) (11, 12). This ECM network not only provides physical scaffolding but also regulates key biological processes, including cell adhesion, proliferation, migration, and differentiation, and acts as a reservoir for signaling molecules such as growth factors (13–15).

In metabolically compromised states, such as obesity or insulin resistance, the ECM becomes fibrotic and inflamed—a condition known as fibro-inflammation (16). While fibro-inflammatory signaling in WAT has been extensively studied and is often attributed to immune–adipocyte interactions during tissue expansion (17–19), much less is known about the physiological role of ECM remodeling during BAT activation. During cold exposure, BAT undergoes rapid expansion to meet thermogenic demand, and this is accompanied by dynamic changes in ECM composition. Recent work has shown that collagen expression is altered as early as 4 hours following cold exposure in mice, and these changes are reversible, suggesting an active, physiological remodeling process that supports thermogenic adaptation (20). However, the homeostatic roles and regulatory mechanisms of this ECM remodeling remain poorly understood. Recently, it was demonstrated that collagen deposition and ECM reorganization impair BAT activity and plasticity in mice subjected to obesogenic conditions (21). Complementary findings in a single-cell transcriptomic study of aging rabbits also revealed extensive ECM remodeling, including increased expression of collagens and ECM-associated genes, in BAT stromal populations (22). These findings suggest that maintaining a permissive and dynamic ECM is essential for sustained BAT activity and that non-adipocyte cells, including vascular and stromal elements, may orchestrate this remodeling by integrating thermogenic cues with matrix homeostasis.

Cells interpret and respond to ECM changes through specialized mechanosensing systems, including integrins (e.g., ITGB1, ITGA5), which anchor cells to ECM ligands and initiate intracellular signaling; focal adhesion complexes (e.g., paxillin, FAK), which link integrins to the actin cytoskeleton; cytoskeletal tension sensors (e.g., talin, vinculin), which convey mechanical force and contribute to cell shape and tension; and mechanosensitive signaling molecules such as RhoA, Rac1, and YAP/TAZ, which regulate gene expression, cytoskeletal dynamics, and cell proliferation in response to mechanical inputs (23). These systems not only detect ECM stiffness and architecture but also actively contribute to ECM remodeling through bidirectional signaling, thereby influencing gene expression, cytoskeletal architecture, and cell fate decisions. In BAT, ECM remodeling is recognized as an essential part of cold-induced thermogenic activation, yet the molecular machinery linking matrix sensing to adaptive thermogenesis remains poorly defined. While much of the mechanotransduction research has concentrated on adipocytes or preadipocytes (23), stromal and vascular cells are also likely to participate in sensing and remodeling the ECM in response to thermal and metabolic cues.

Among known mechanosensory systems, G protein–coupled receptors (GPCRs), and particularly the adhesion GPCR (aGPCR) family, have garnered interest due to their large extracellular domains capable of binding matrix ligands and mediating mechanical signal transduction (24). aGPCRs are structurally equipped to function as mechanosensors, and several members have been shown to mediate mechanical signal transduction in response to ECM cues or shear stress with metabolic consequences (25, 26). For example, Adgrg6(Gpr126), a prototypic mechanosensor regulates Schwann cell development via ECM interaction (27, 28) and loss of its expression in chondrocytes and tendon cells leads to spine defects (scoliosis), due to impaired mechanosensing and altered tissue stiffness (29); Adgrg1(Gpr56) controls myoblast fusion (30) and muslce hypertophy via interraction with collagen type III (31) and white adipocyte differentiation, particularly acting on adipocyte progenitors and profibrotic fibroblasts (32). Adgrl2(Latrophilin) is a bona fide mechanosensor for shear forces, and it translates the mechanical stimulus of blood flow into biochemical signals that drive endothelial cell alignment, new vessel formation, and structural remodeling of arteries to meet the hemodynamic demands (33). In addition, other aGPCRs have been linked with adipogenesis, such as Adgrg2(Gpr64) (34) and thermogenesis, such as Adgra3 (35), although it is not clear whether these effects involve mechanosensing.

Although the mechanosensing roles of aGPCRs in thermogenic tissue are yet to be fully elucidated, their structural properties and emerging metabolic functions position them as compelling candidates for linking ECM dynamics to thermogenic remodeling. Notably, the expression profiles and cell-type specificity of many aGPCRs in BAT remain poorly characterized. In this study, we aimed to map aGPCR expression in BAT using single-nuclei transcriptomic analysis. We found that aGPCRs are the second most abundant GPCRs in BAT, after Class A receptors. This approach revealed Adgrf5(Gpr116) as one of the most highly enriched aGPCRs, with selective expression in endothelial cells in both mouse and human BAT. Through a combination of genetic deletion models and cold exposure challenges, we examined the role of Adgrf5 (Gpr116) in thermogenic remodeling, vascular adaptation, and matrix homeostasis. Our findings show that loss of endothelial Adgrf5(Gpr116) leads to endothelial-to- mesenchymal transition (EndMT) and fibro-inflammatory transcriptional reprogramming during cold-induced thermogenic stress, ultimately impairing BAT function. These results reveal a previously unrecognized endothelial mechanism by which Adgrf5(Gpr116) regulates BAT tissue remodeling and thermogenic response.

## 2. Methods and Materials

### 2.1 Animal models

All mice (Mus musculus) were born and maintained in a temperature-controlled (22 °C) room on a 12-h light/dark cycle. Mice were fed with a regular unrestricted chow diet (Kliba Nafag #3437, Provimi Kliba AG, Kaiseraugust). Chow-fed mice were housed 4– 5 mice per cage. Mice were singly caged for the duration of the indirect calorimetry experiments. Body weight was measured with a table scale, and body composition (lean and fat mass) was measured via nuclear magnetic resonance technology (EchoMRI). All animal handling procedures were performed under the European Union directives and the German Animal Welfare Act and have been approved by the institutional animal welfare officer and local authorities under animal licenses (ROB- 55.2-2532.Vet_02-21-133).

#### Transgenic mouse models

*ADGRF5(GPR116) knockout KO*: *ADGRF5(GPR116)* knockout (KO) mice have been previously described by Yang et al. (36) and Fukuzawa T et al. (37) which carry a deletion of exon 2, which encodes for the start codon and signal peptide (SP) of the *Adgrf5(Gpr116)* gene. The mice were a present from Brad Croix. They were backcrossed ten times, already upon arrival, and they were on a clean C57Bl6N genetic background. Genotyping was done with primers: WT: GGAGGCTCTGTGCGTTTC, R1: CTGTGGACATGATGAAGGGTG, R2: CTCCCTGAATCATAGTCTAGTCTCC. PCR reaction was run with DreamTaq (ThermoScientific) using the following PCR protocol: Denaturation 95°C (5min), (denaturation 95°C 30sec, annealing: 60°C 30sec, elongation: 72°C 1min) for 35 x cycles and final extension 72°C for 10min.

*Ucp1CrexADGRF5(GRP116)f/f*: For conditional knockdown mouse lines, ADGRF5(GPR116) flof/flox, with flox sequences surrounding exon2, on a C57Bl6N genetic background, B6N.Cg-Adgrf5tm1.1Bstc/J, Strain #:022505 from the Jackshon Lab. were crossed with Ucp1Cre (C57BL6J genetic background, B6.FVB-Tg(Ucp1- cre)1Evdr/J, strain 024670 Jaxon Labs), to generate Ucp1CreGPR116f/f mice. PCR genotyping was done with primers: For Ucp1Cre forward 1: CAAGGGGCTATATAGATCTCCC, Cre reverse 1: ATCAGAGGTGGCATCCACAGGG, Cre reverse 2: GTTCTTCAGCCAATCCAAGGG, and for Adgrf5(Gpr116)flox/flox the same primers as in the genotyping of the ADGRF5(GPCR116) KO mice were used. PCR was run with DreamTaq (ThermoScientific) with the following protocol: Denaturation 95°C (5min), (denaturation 95°C 30sec, annealing: 60°C 30sec, elongation: 72°C 1min) for 35 x cycles and final extension 72°C for 10min. Results were assesed on an agarose gel (2%) based on band sizes: UCP1Cre band 336bp, WT band: 554bp. For ADGRF5(GPR116)flox/flox band 340bp and WT band 292bp.

*Cdh5CreERT2xADGRF5(GRP116)f/f*: C57BL/6- Tg(Cdh5-cre/ERT2)1Ra were generated by Ralf H Adams (38) , and they were imported from the Mary Lyon Center at MRC Harwell (EMMA ID: EM 14891). Genotyping for Cdh5CreERT2 was done with primers Forward: TCCTGATGGTGCCTATCCTC and Reverse: CCTGTTTTGCACGTTCACCG usign PCR DreamTaq (ThermoScientific) and the following protocol: Initial denaturation 95oc 1min, denaturation 95°C 10sec, annealing 60°C 10 sec, elongation 72°C 1min foro 29 cycles, final extension 72°C 30sec. Results were assessed by agarose gel and Cdh5Cre positive band was at 594bp.

The Cdh5Cre expression was induced by intraperitoneal (i.p.) injections of tamoxifen (Sigma, T5648) at 100 mg/kg/day for 5 consecutive days. Tamoxifen was prepared as a 20 mg/mL working solution by dissolving 60 mg in 0.5 mL pure ethanol, followed by the addition of 2.5 mL corn oil. Mice were allowed a 2-week washout period post- injection to ensure complete recombination and clearance of tamoxifen activity. Recombination success was checked with qPCR reaction on Cd31+ ΒΑΤ isolated cells, with qPCR primers against *Adgrf5(Gpr116)* FW: TTCTTCAGAAGCTGCGCTGA and RV: CCCGGCTAGCTCATGTTCTT.

### 2.2 Isolation of BAT endothelial cells

Endothelial cells were isolated from BAT using magnetic-activated cell sorting (MACS) following an adapted protocol from Joerg Heeren lab at UKE Hamburg(39). Culture dishes were pre-coated with 1.5% gelatin and 10 μg/mL human fibronectin. Cells were maintained in Lonza EGM-2MV basal medium supplemented with 10% FCS, 0.1 mg/mL kanamycin, and 100 ng/mL VEGF, with daily media changes.

Fat pads were aseptically harvested and placed in 0.9% NaCl on ice. Tissues were minced and digested in isolation buffer (123 mM NaCl, 5 mM KCl, 1.3 mM CaCl₂, 5 mM glucose, 100 mM HEPES pH 7.4) supplemented with 600 U/mL collagenase II and 1.5% BSA at 37°C for 40 minutes BAT with agitation. The digested tissue was filtered through 100 μm and 40 μm cell strainers and centrifuged at 600 × g for 5 minutes at 4°C.

Endothelial cells were purified using a two-step MACS protocol. First, CD11b⁺ cells (macrophages) were depleted by negative selection using CD11b MicroBeads (Miltenyi Biotec) and LS columns. The CD11b⁻ fraction was then incubated with CD31 MicroBeads and Isolectin B4 FITC followed by anti-FITC MicroBeads for positive selection of endothelial cells. CD31⁺ endothelial cells were eluted from LS columns and seeded in Lonza-EC medium for culture.

### 2.3 Cold challenge, adrenergic stimulation, and indirect calorimetry

Energy expenditure (EE), locomotor activity, Respiratory exchange ratio (RER), and food intake were measured and calculated by combined indirect calorimetry over indicated periods (PhenoMaster; TSE Systems, Bad Homburg vor der Höhe, Germany) as described previously (40). Before any measurements were collected, mice were single-caged and acclimated in the PhenoMaster: TSE system for 7 days. During the acute cold exposure challenge, the temperature was dropped from 22 °C to 4 °C within 30 minutes, and measurements were collected for up to 8 hours (h), when mice were sacrificed or returned to 22 °C. For the prolonged cold challenge protocols, the ambient temperature was gradually dropped from 22 °C to 8 °C over 7 days, being reduced by 2°C every 12h. Indirect calorimetry data were tabulated using the CalR tool (41). For pharmacological activation of the Adrb3 (β3 adrenergic receptor) signaling, mice were injected i.p. with 2mg/kg of CL 316243 (Sigma, C5976).

### 2.3 Core body temperature measurements

Core body temperature was measured with a telemetric probe (TA-F10, Data Sciences International, Cat# TA-F10), which was surgically implanted into the peritoneal cavity of mice under aseptic conditions. Anesthesia was induced using 2.5% isoflurane. Before surgery, mice received a single subcutaneous dose of buprenorphine (0.1 mg/kg). Additionally, carprofen (20mg/kg, s.c.) was administered after surgery and at least once daily for two days postoperatively. No surgical complications were observed, and mice were allowed a minimum recovery period of one week before data collection commenced. During telemetry data recording, mice remained in their home cages, which were positioned on telemetry receivers (RPC-1, Data Sciences International). Data acquisition was carried out using an MX2 matrix (Data Sciences International), with continuous recording via PhysioTel software in PONEMAH Physiology Platform (PhysioTel system and open the software (PONEMAH Physiology Platform v.6.30)). The collected data were subsequently organized into 60-minute intervals using Microsoft Excel.

### 2.4 Histology and microscopy

Standard Formalin-fixed paraffin-embedded (FFPE) sample preparation has been performed on BAT and iWAT. General characteristics of tissue structure were assessed by H and E staining (H&E staining).

For 2D-CD31 staining, stained sections were scanned with an AxioScan 7 digital slide scanner (Zeiss) equipped with a ×20 magnification objective. Tissue detection with focus points was applied automatically. Digitized slides were loaded into the Visiopharm image analysis system, and the algorithm was used to classify the CD31- marked area and its staining intensity.

For 3D-whole mount imaging of vasculature staining of adipose tissue, iWAT was dissected and fixed overnight at 4°C in 4% paraformaldehyde (PFA). After washing the tissue thrice in PBS it was embedded in 3% agarose (in H_2_O) to aid with cutting sections. Agarose embedded iWAT was sliced into 200 µm sections using a fully automated vibrating blade microtome Leica VT1200S (Leica Biosystems). Tissue sections were collected in blocking solution containing 1% (w/v) bovine serum albumin (BSA) and 1% (v/v) Triton^TM^X-100 in PBS and incubated for 2 h at RT. Following fixation and blocking, primary antibodies were added diluted in blocking solution. Tissue sections were incubated overnight at 4°C. After washing thrice in PBS, secondary antibodies, diluted in blocking solution containing DAPI were added. Tissue sections were incubated overnight at 4°C protected from light. After washing thrice in PBS sections were mounted to a microscopy slide using Aqua-Poly/Mount (Fisher Scientific). BAT sections were recorded in confocal z-stacks at the Microscopy Core Facility of the Medical Faculty at the University of Bonn (Leica SP8).

Antibodies : CD31 BD Pharmingen Cat. No 553370, PLIN1 Abcam Cat. No ab61682.

### 2.5 Immunobloting

Frozen tissue was homogenized in Lysis buffer (10mM Tris-HCl, pH 8.0, 1mM EDTA, 0.5mM EGTA, 1% Triton X-100, 0.1% Sodium Deoxycholate, 0.1% SDS, 140mM NaCl, protease inhibitors and phosphatase inhibitors (ThermoFisher Scientific). Adipose tissue lysates were further centrifuged to remove any adipocyte lipid layer present. Protein concentrations were determined using a bicinchoninic acid assay (BCA) kit (ThermoFisher Scientific). Proteins were resolved on 4-12% bis-Tris gels (Invitrogen), then transferred to PVDF membranes (Bio-Rad). For tyrosine hydroxylase we used Tyrosine Hydroxylase, clone LNC1 from Sigma-Aldrich catalogue no. MAB318. Loading control was VCP Abcam ab11433.

### 2.6 ELISA Kits

For measurement of serum thyroid hormone levels (T3) we used the mouse Triiodothyronine (T3) ELISA Kit from BIOZOL, cat. Number MBS262762-96.

### 2.7 Serum analyser

Measurements of serum lipids and lactate were performed using the serum analyzer (AU480 Beckman Coulter).

### 2.8 Measurement of norepinephrine in mouse plasma

Plasma norepinephrine was quantified using high-performance liquid chromatography (HPLC) coupled with electrochemical detection (EcD), as previously described (42). Sample preparation followed the protocol provided by RECIPE Chemicals + Instruments GmbH for catecholamine extraction in human plasma with the ClinRep® complete kit, which includes all necessary reagents and materials for the extraction of the desired analytes. Due to the limited volume of mouse plasma/serum, modifications to the standard protocol were required. Specifically, 50 µL of mouse plasma/serum was mixed with 100 µL of 0.3 M perchloric acid (in water) and 5 µL of internal standard (DHBA, 1 ng/µL). After thorough mixing, the sample was centrifuged at 9,000 rpm for 5 minutes. The resulting supernatant was charged to the sample preparation column, which was then shaken for 10 minutes. Solvent was removed using a vacuum manifold, and the column was washed three times with 1 mL of wash solution to eliminate interfering substances. After drying the column, 150 µL of elution reagent was added. The column was shaken for 5 min, and then the catecholamines were eluted from the extraction column via centrifugation. A 20 µL aliquot of the final eluate was injected into the HPLC-EcD system for analysis.

### 2.9 Nuclei isolation and fixation

Brown adipose tissue (BAT) from two mice per ambient temperature condition was pooled for each sample. All steps were performed on ice using pre-cooled reagents, tubes, and a swing-bucket centrifuge to preserve nuclear integrity. Snap-frozen BAT tissue was first fragmented into small pieces using a mortar and pestle pre-cooled with liquid nitrogen. The frozen tissue fragments were immediately transferred into 50 mL gentleMACS™ C Tubes (Miltenyi Biotec, 130-128-024) prefilled with nuclei extraction buffer (Miltenyi Biotec) at a ratio of 2 mL per 200 mg of tissue, supplemented with 10,000 units of RNasin® Plus RNase Inhibitor (Promega, N2615).

Tissue homogenization was carried out using the gentleMACS™ Dissociator device, running a 5-minute program optimized for nuclei extraction. The program was repeated twice to ensure complete dissociation of the tissue. The resulting homogenate (typically 2–4 mL per sample) was brought up to a final volume of 10 mL using resuspension buffer consisting of 1% BSA, 2 mM MgCl₂, and 0.2 U/μL RNase inhibitor. The homogenate was then filtered through a 70 μm cell strainer (Miltenyi Biotec, 130-098-462) into a fresh 15 mL conical tube and centrifuged at 500 × g for 5 minutes at 4 °C. The supernatant was removed using a vacuum pump, and the pellet was resuspended in 1 mL of resuspension buffer, filled again to 10 mL, and centrifuged under the same conditions to perform a second wash. The final pellet was resuspended in 500 μL of resuspension buffer, filtered through a 40 μm cell strainer, and transferred to a 1.5 mL low-binding microcentrifuge tube.

Nuclei were counted manually using a hemocytometer (Sigma-Aldrich, BR717805- 1EA) and subsequently fixed according to the Parse Biosciences Nuclei Fixation Protocol v2. Following fixation, each nuclei suspension was counted again and divided into two aliquots: one for pre-barcoding quantification and one reserved for the barcoding step. Fixed nuclei were stored at –80 °C until further processing.

### 2.10 Parse Biosciences single nuclei library preparation and sequencing

Fixed nuclei suspensions from each experimental condition were processed for single-nucleus transcriptomic profiling using the Split-Seq combinatorial barcoding method, as implemented in the Parse Biosciences Evercode™ Whole Transcriptome v2 kit. For each condition, we targeted the recovery of approximately 25,000 to 30,000 barcoded nuclei.

Following the barcoding steps, cDNA concentration from each library was measured using the Qubit High Sensitivity DNA Kit (Life Technologies, Q33230). The fragment size distribution of the cDNA was assessed with the Agilent Bioanalyzer using the High Sensitivity DNA Kit (Agilent Technologies, 5067-4626).

Library preparation steps, including cDNA fragmentation and Illumina adaptor and index ligation, were performed according to the manufacturer’s instructions provided with the Parse Biosciences kit. Final libraries were submitted to the Helmholtz Munich Genomics Core Facility for sequencing on an Illumina NovaSeq X Plus system.

### 2.11 Single-nuclei RNA sequencing data analysis

The raw fastq files generated for each sub-library were processed using the demultiplexing pipeline provided by Parse Biosciences (v2, Parse Biosciences, Inc., 2024). The reference genome utilized for the alignment was the GRCm39. Subsequently, the 8 sub-libraries were merged using a function that combines the data of all the sub-libraries, which is included in the same version of the split-seq pipeline. The count matrices derived from the combined analysis of the 8 sub-libraries for each sample were then used as input in R programming language and more specifically the Seurat package v5 (43).

The count matrices of the different conditions were merged using the *merge()* function of the Seurat library to generate a Seurat object by keeping the condition information. The data was subset by keeping the nuclei that have less than 5,000 features, 20,000 counts, and 15% mitochondrial gene content. The filters aim to reduce the information that comes from low-quality nuclei, doublets, and ambient RNA. The high-quality nuclei gene expression counts were scaled using the ScaleData Seurat function, and after the pre-processing, the standard Seurat analysis, including principal component analysis (PCA), FindNeighbors, and finally FindClusters. Additionally, the integration tool used was Harmony (44), which was combined with the Leiden algorithm (45) (resolution 0.25) for the cluster identification. After pre-processing a joint embedding, Uniform Manifold Approximation and Projection (UMAP) was applied for visualisation of the different cell populations for BAT.

The Seurat FindMarkers function with the MAST package (46) differential gene expression method was applied for identifying gene markers that are uniquely highly expressed for each cluster. In order to speed up the FindMarkers() function for each cluster, we have used a bash script for parallelization using multiple nodes on the high-performance computing cluster (HPC). Based on the gene markers, literature, and recent single cell/nuclei data we were able to annotate the clusters and identify the different tissue cell types. Finally, the FindMarkers function with the MAST package (46) was applied for identifying condition and genotype-dependent differentially expressed genes.

### 2.12 GPCRs expression analysis

The databases IUPHAR/BPS Guide PHARMACOLOGY (www.guidetopharmacology.org) and the HUGO Genome Nomenclature Committee (www.genenames.org) were utilized assing GPCRs to classes ( Class A, Class B, Frizzled and Adhesion GPCRs). The R library scCustomize (Marsh SE,2021) and more specifically the function Percent_expressing() (Marsh SE,2021) was applied for calculating the percentage of GPCR classes per condition. We considered relevant GPCRs which expressed in more than 1% of nuclei in a cell population (cutoff of receptor frequency 1%).

The same GPCR expression analysis was performed using a publicly available single-nuclei human BAT dataset (47). We visualized the common and unique GPCR receptors in mouse cold-exposed conditions or room temperature (RT), as well as in the human dataset with upset plots from the UpsetR package (48).

### 2.13 Gene Ontology analysis

The GSEA methodology and the clusterprofiler (49) were applied for gene ontology analysis. Differentially expressed genes for each condition and genotype comparison were combined in one list, which was used as an input in the compareCluster() (49) function with pvalueCutoff 0.05. Additionally, the function simplify() (49) was applied in the compareCluster() result for reducing the redundant gene ontologies.

### 2.14 Cell-cell interaction analysis in BAT

The CellChat (50) analysis was applied to calculate intercellular interactions. The analysis was performed for each condition (WT_22°C, WT_8°C, KO_22°C, KO_8°C) separately. Afterwards, the individual CellChat objects were merged with the mergecellchat() function, to summarize which cell-cell interactions showed condition- specific changes and which were common between conditions.

### 2.15 Regulons analysis

We performed transcription factor regulon analysis using the decoupler (51) library and the DoRothEA gene regulatory network (52) , which contains information for transcription factors and their target genes from mouse and human. The transcription factor analysis was performed for the subset of arterial and capillary endothelial for each condition in the BAT (WT_22°C, WT_8°C, KO_22°C, KO_8°C) separately. The results of the analysis for each condition were visualized with the pheatmap heatmap library.

### 2.16 Software and algorithms details, please see Supplementary Table 1

### 2.17 Statistics and graph generation

Energy expenditure was calculated using the Weir equation. Data analysis and statistics (which include ANOVA and ANCOVA) were performed using GraphPad Prism. The CalR platform was used to analyze the indirect calorimetry data.

### 2.18 Data availability

This paper does not report original code. Web application app for data exploration can be found and the link: https://shiny.iaas.uni-bonn.de/Shiny-GPCR/ (username: first_glance , password: 3LNRearr7Xdv). The raw files and analysed objects can be shared upon request to the corresponding author.

### 2.19 Use of AI tools

We used OpenAI’s ChatGPT (2024 version) for troubleshooting and optimizing code during the analysis of single-nucleus transcriptomic data, as well as for improving clarity, grammar, and structure throughout the text. All scientific content, data interpretation, and final wording were generated and approved by the authors.

## 3. Results

### 3.1 Single-nuclei RNA sequencing screen identifies adhesion GPCRs as the second most abundant GPCR class in BAT

Previous efforts to screen for GPCRs in BAT have been performed via pre-designed primers for qPCR or multiplexing arrays, on either whole mouse BAT tissue (53, 54) or in vitro differentiated brown adipocytes (55). However, a single-cell level unbiased screen is missing. To describe the distribution of GPCRs in different cell types in mouse and human BAT, we utilized single-nuclei RNA sequencing from mouse and human BAT. To further examine the effect of cold exposure on GPCR expression in different cell types, we utilised BAT tissue from mice housed at 22°C and mice housed at 8°C for 2 weeks. We obtained 44,635 high-quality nuclei from mouse BAT (Fig. 1A). For single-nucleus GPCR analysis in human BAT, we used a publicly available dataset (47). The human BAT dataset contained 36,590 nuclei (Fig. 1B). Cell clustering identified 12 distinct cell types in mouse BAT, spanning major populations including brown (BAd) and white (WAd) adipocytes, adipogenic progenitors (APCs) and more fibroblast like proliferating progenitors (Prol.prog.-Fibro.), endothelial cells (arterial-ArtECs, capillary-CapECs, lymphatic-LECs) and mural cells (MuralC) containing vascular smooth muscle cells and pericytes, immune cells; specifically macrophages (Macro), myogenic progenitors (MyoProg.), myocytes (MyoC) and schwann cells (SC), which we annotated based on canonical gene markers (Suppl. Fig. 1A). For the human dataset we identified in common to the mouse BAT, brown adipocytes (BAd), fibroblast like proliferating progenitors (Prol.prog-Fibro.), myocytes (MyoC), endothelial cells (ArtECs/CapECs), macrophages (Macro) and in addition, human BAT contained preadipocytes (Pread.) and a big number of immune cells; T-cells, class-switched memory B cells, dendritic cells (Fig. 1B and Suppl. Fig. 1B).

**Figure 1:**
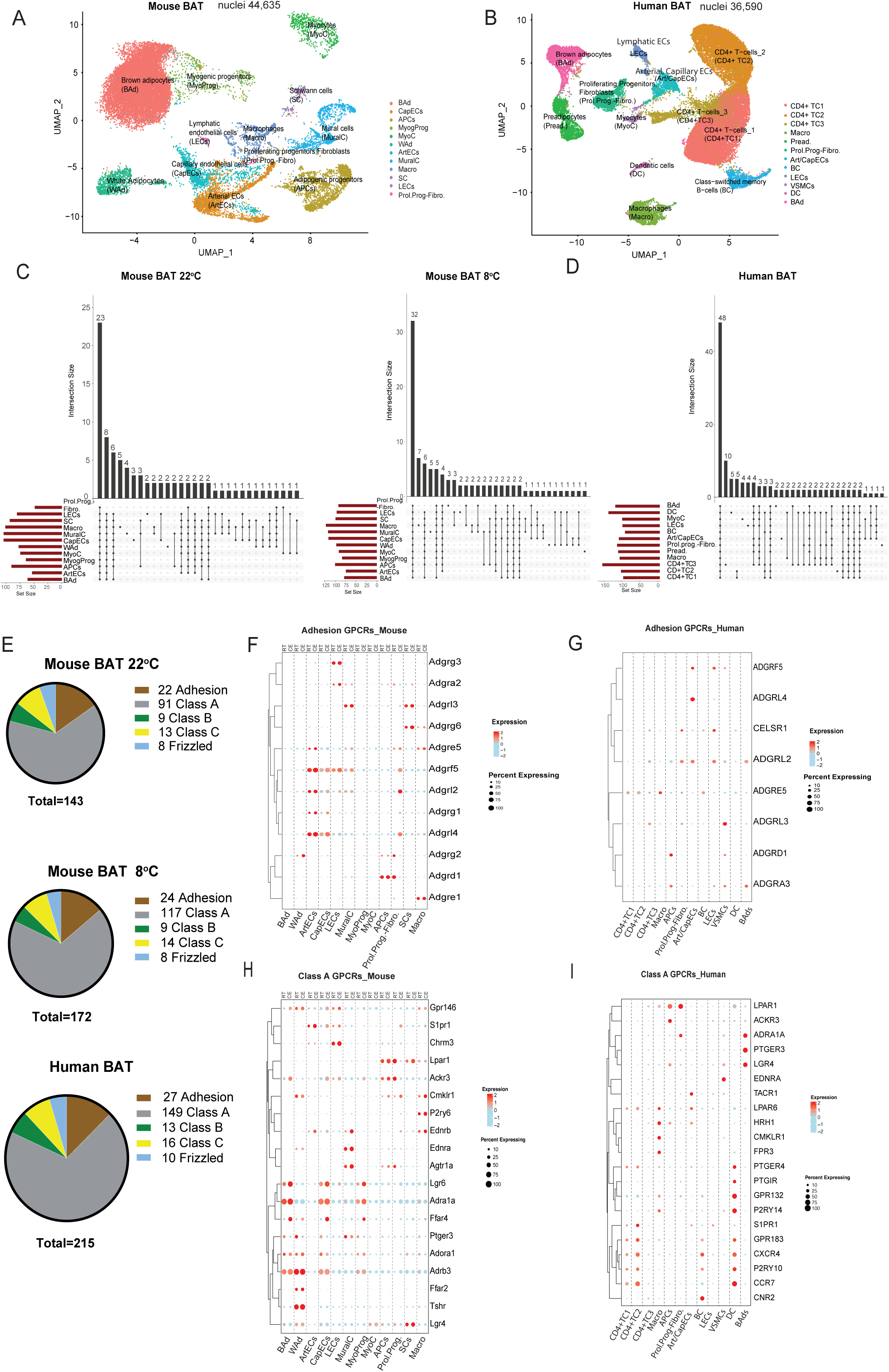
Single-nuclei analysis of GPCRs in mouse and human BAT. UMAP plots showing clustering of nuclei from (A) mouse BAT, (B) human BAT. (C-D) Upset plots showing the number of common and unique GPCRs across different cell types for mouse BAT (C) and human BAT (D). (E) Pie charts showing the percentage of represented classes of GPCRs in mouse and human single-nuclei datasets. (F-I) Dot plot expression of indicated GPCRs in mouse and human BAT cell types. Mice were either kept single housed at 22 °C or 8°C for 14 days. Single-nuclei RNAseq is a pool of BAT from two mice per ambient temperature condition. Mice were males, C57Bl6N, 10-12 weeks old.

We detected 143 GPCRs in mouse BAT at room temperature (22 °C), 172 in mouse BAT after 2 weeks of cold exposure (8 °C), and 215 in human BAT (Extended File 1). Most of the detected GPCRs were not uniquely expressed in a single cell type. Notably, in mouse BAT several GPCRs showed macrophage-specific expression at 22 °C (e.g., *Cnr2, Gpr141, Gpr183, C5ar1, Adgre4*) and at 8 °C (Cnr2, Cxcr5, Gpr171, Gpr183, C3ar1) (Fig. 1C and Extended File 2). In human BAT, immune CD4+ T cells exhibited the highest number of cell–type–specific GPCRs, including *CHRM2, GPR139, GPR149, LPAR3, RXFP2, GHRHR, GRM4, and GPRC5D* (Fig. 1D and Extended File 2).

In addition, we compared our single-nuclei datasets with previously published data on detected GPCRs in total mouse BAT (54) and mouse brown adipocytes and preadipocytes (55). Although some GPCRs expressed at very low levels—such as *Gpr3*—were not detected in our dataset, we recovered a broad range of GPCRs that were previously reported in mouse BAT, preadipocytes, and differentiated brown adipocytes (Extended File 3). The overall high coverage of GPCRs in our dataset, including many that have been reported in prior qPCR or bulk RNA-seq studies, demonstrates the sensitivity and specificity of our single-nucleus RNA-seq strategy.

We classified the detected GPCRs according to standard GPCR family nomenclature into Class A, Class B, Class C, Frizzled, and Adhesion GPCRs (aGPCRs). The majority of GPCRs detected in both mouse and human BAT belonged to Class A, followed by aGPCRs, as the second most represented family (Fig. 1E).

Looking at the top-expressed adhesion GPCRs (aGPCRs), we observed a strong cross-species correspondence between mouse and human BAT, which was not evident for class A GPCRs (Fig. 1F-1I), suggesting a potentially conserved role for aGPCRs in thermogenic fat. Among the identified aGPCRs, we detected *Adgra3* in adipocytes, previously reported to regulate brown adipocyte thermogenesis((35), and *Adgrd1* in adipocyte progenitor cells (APCs) and proliferating progenitor–fibroblast populations, consistent with its established role in controlling adipogenesis in white adipose tissue (56). Notably, *Adgrf5* and *Adgrl4* emerged as the two most highly expressed aGPCRs in both human and mouse BAT, with clear enrichment in vascular endothelial cells across species (Fig. 1F, 1G). While *Adgrl4* has a well-characterized function in angiogenesis and vascular integrity((57), *Adgrf5(Gpr116)* has been implicated in endothelial homeostasis in the lung (58) and the brain((59), but its role in adipose tissue vascular biology and thermogenic adaptation has not been explored. Given the central role of the endothelium in regulating tissue remodeling and function, we focused on investigating the contribution of *Adgrf5(Gpr116)* to BAT adaptation under thermogenic stress.

### 3.2 Cold exposure induces aGPCR-mediated cell-to-cell contact interactions between BAT non-adipocyte cells

Cold exposure stimulates angiogenesis, which enhances nutrient delivery to active BAT (60). Beyond this classical vascular role, endothelial cells also contribute to tissue remodeling through dynamic intercellular communication (61). To investigate how cold exposure shapes cellular cross-talk in BAT—particularly involving vascular cells—we applied the CellChat algorithm, which infers cell–cell communication networks based on ligand–receptor expression across defined cell types.

Analysis of the single-nucleus BAT dataset revealed that cold exposure substantially increased the total number of predicted interactions between all cell types (Fig. 2A). However, vascular cells (endothelial and mural cells) emerged as central communication hubs, displaying high levels of interaction both within the vascular compartment and with other cell types, especially adipogenic progenitors (APCs) and proliferating progenitor-fibroblast-like cells (Fig. 2A). Specifically for ECs these interactions were dominated by cell–cell contact, ECM–receptor, and secreted signaling categories, with a notable increase in all three upon cold exposure, while non-protein-based signaling played a minor role (Fig. 2B).

**Figure 2:**
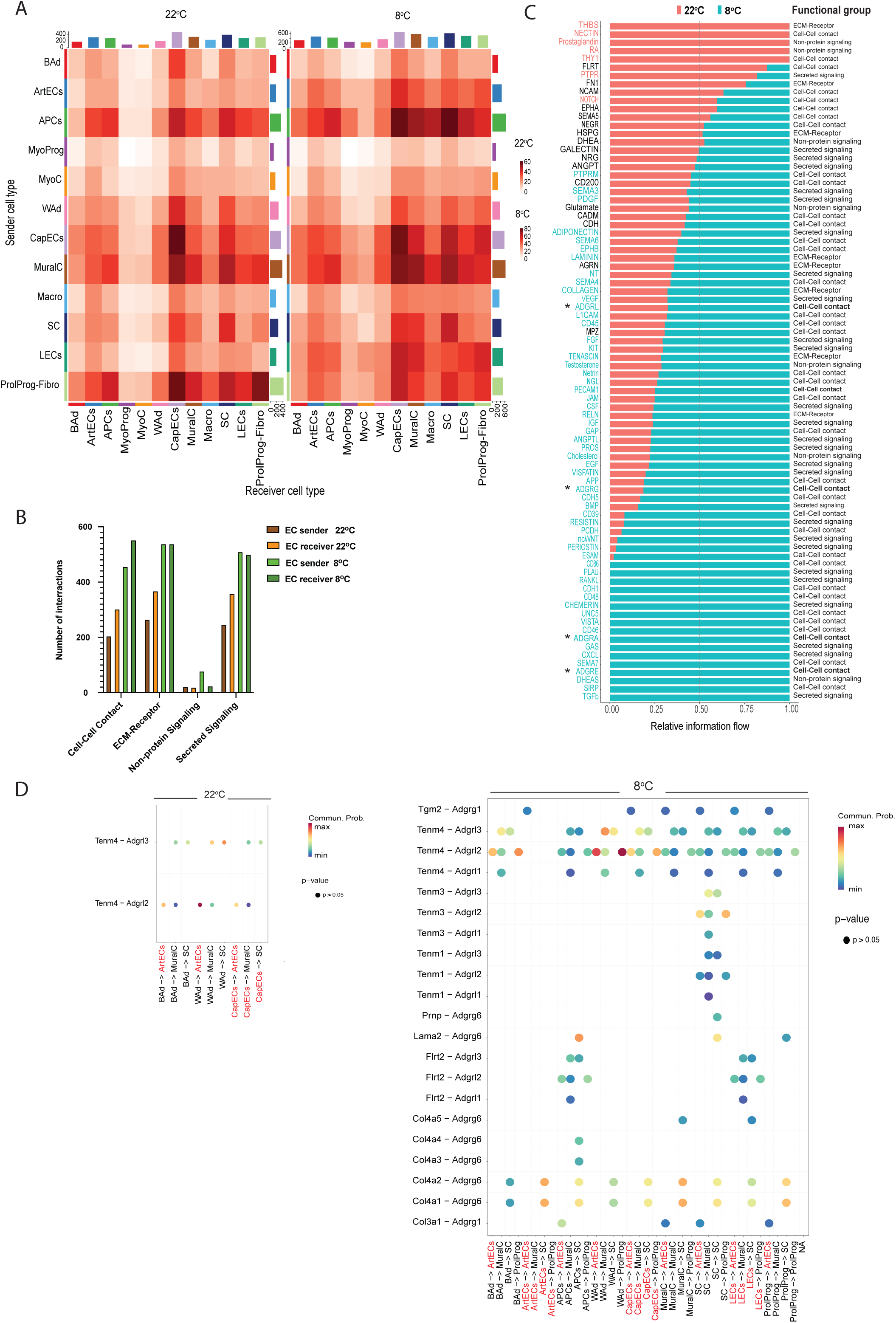
Temperature-related interactions between cell types in BAT. (A) Heatmaps showing the interaction score between indicating cell types, as senders or receivers (B) Number of interactions of different categories (cell-cell contact, ECM-receptor, non-protein signaling, and secreted signaling involving endothelial cell (EC) as sender or receiver at indicated ambient temperatures. (C) Bar plots showing the relative information flow for significant communication pathways between all BAT cell types.* indicates the aGPCRs pathways. (D) Dot plots presenting significant ligand- aGPCRs receptors predicted to participate in the communication of indicated cell types. The colours of the circles correspond to communication probability. ECs are highlighted in red colour.

Given the earlier finding that aGPCRs are among the most enriched GPCRs in BAT and are prominently expressed in vascular cell populations, we next examined whether aGPCR signaling contributes to the increased cold-induced communication landscape. Indeed, several predicted cold-specific interactions involved aGPCR family members—including ADGRL, ADGRG, ADGRA, and ADGRE receptors—engaged in cell–cell contact signaling (Fig. 2C). These pathways were particularly enriched in non- adipocyte lineages and frequently involved ECs, and ECM components such as collagens serving as ligands. These findings support a model in which aGPCRs act as sensors of mechanical and matrix-derived cues and play a key role in mediating cold-induced cellular crosstalk between the vascular and stromal compartments of BAT (Fig. 2D).

### 3.3 Global deletion of Adgrf5(Gpr116) distrubs thermoregulation upon prolonged cold exposure

Among the aGPCRs enriched in the BAT vasculature, Adgrf5(Gpr116) emerged as a highly expressed receptor in ECs of both mouse and human BAT. Our CellChat analysis further suggested that aGPCRs may mediate enhanced intercellular signaling in response to cold exposure, particularly between vascular and progenitor compartments. However, the functional contribution of Adgrf5(Gpr116) to BAT remodeling and thermogenic adaptation has not been characterized to date.

Publicly available datasets confirmed that *Adgrf5(Gpr116)* gene is a pan-endothelial receptor (Suppl. Fig. 2A) and we measured its expression primarily in the lung, followed by BAT, heart, and skeletal muscle (Suppl. Fig. 2B). Notably, among major metabolic tissues, *Adgrf5(Gpr116)* was selectively induced in BAT after cold exposure, but not in WAT, liver, or skeletal muscle (Suppl. Fig. 2C). Within BAT, its expression is predominant in all EC populations, with lower expression detected in a subset of muscle cells, brown adipocytes, preadipocytes, and mural cells (Suppl. Fig. 2D).

Several mouse models have been developed to delete Adgrf5(Gpr116), a gene spanning 29 exons. One commonly used model involves the deletion of exon 2, which encodes the signal peptide and the translation start site, rendering the protein non-functional (36). This exon 2 deletion results in a progressive pulmonary phenotype characterized by late-onset alveolar dysfunction and emphysema, beginning around 18 weeks of age, phenocopying the phenotype observed in endothelial-specific Adgrf5(Gpr116) knockouts using Tie2-Cre (59). Similarly, a broader deletion spanning exons 4–21, which removes the majority of the extracellular, transmembrane, and cytoplasmic domains, reproduces the same pulmonary phenotype (59). These consistent findings across multiple models indicate that Adgrf5(Gpr116) is required for maintaining pulmonary homeostasis and that its physiological functions are, to a great extent, mediated through endothelial expression. Thus, the phenotype observed in global knockout models possibly reflects loss of *Adgrf5(Gpr116)* function in ECs.

To determine whether *Adgrf5(Gpr116*) plays a physiological role in cold-induced thermogenesis, we next utilized a global *Adgrf5(Gpr116*) knockout mouse model (exon2 deletion) (36) and subjected these mice to acute and prolonged cold exposure. During an acute cold challenge (4°C for 8 hours), both genotypes exhibited comparable increases in energy expenditure (EE), oxygen consumption (VO₂), carbon dioxide production (VCO₂), but interestingly they showed lower respiratory exchange ratio (RER) and a trend for lower food intake and locomotor activity, compared to WT controls (Fig. 3A–F). In contrast, under prolonged cold exposure (8°C for 14 days), ADGRF5(GPR116) KO mice showed clear signs of impaired cold adaptation. Specifically, they exhibited significantly lower EE, VO₂, and VCO₂ during the dark (active) phase compared to WT controls, while RER remained unchnanged during the dark phase and sliglty increased in the ADGRF5(GPR116) KO mice during the light phase, food intake was slightly lower in the ADGRF5(GPR116) KO during the dark phase, whereas activity was dramatically lower in the dark phase (Fig. 3G-3M). Most notably, core body temperature (Tₘ) was markedly lower in ADGRF5(GPR116) KO mice during both the dark and light phases (Fig. 3N). These differences appeared from the first days of cold and they were maintained over 14 days, without leading to lethality. In addition, we report here that we observed exactly the same phenotype in ADGRF5(GPR116) KO females, supporting no sex dependent responses to cold. Of note, under standard housing conditions (22 °C), we observed no differences in EE, VO2, VCO2, RER, food intake, locomotor activity (Suppl. Fig. 3A-3G), pointing to a cold challenge-related underlying reason for the differences observed at 4 °C. Heart and spleen weight were higher though in the ADGRF5(GPR116) KO compared to the WT controls, pointing to systemic complications in this mouse model.

**Figure 3:**
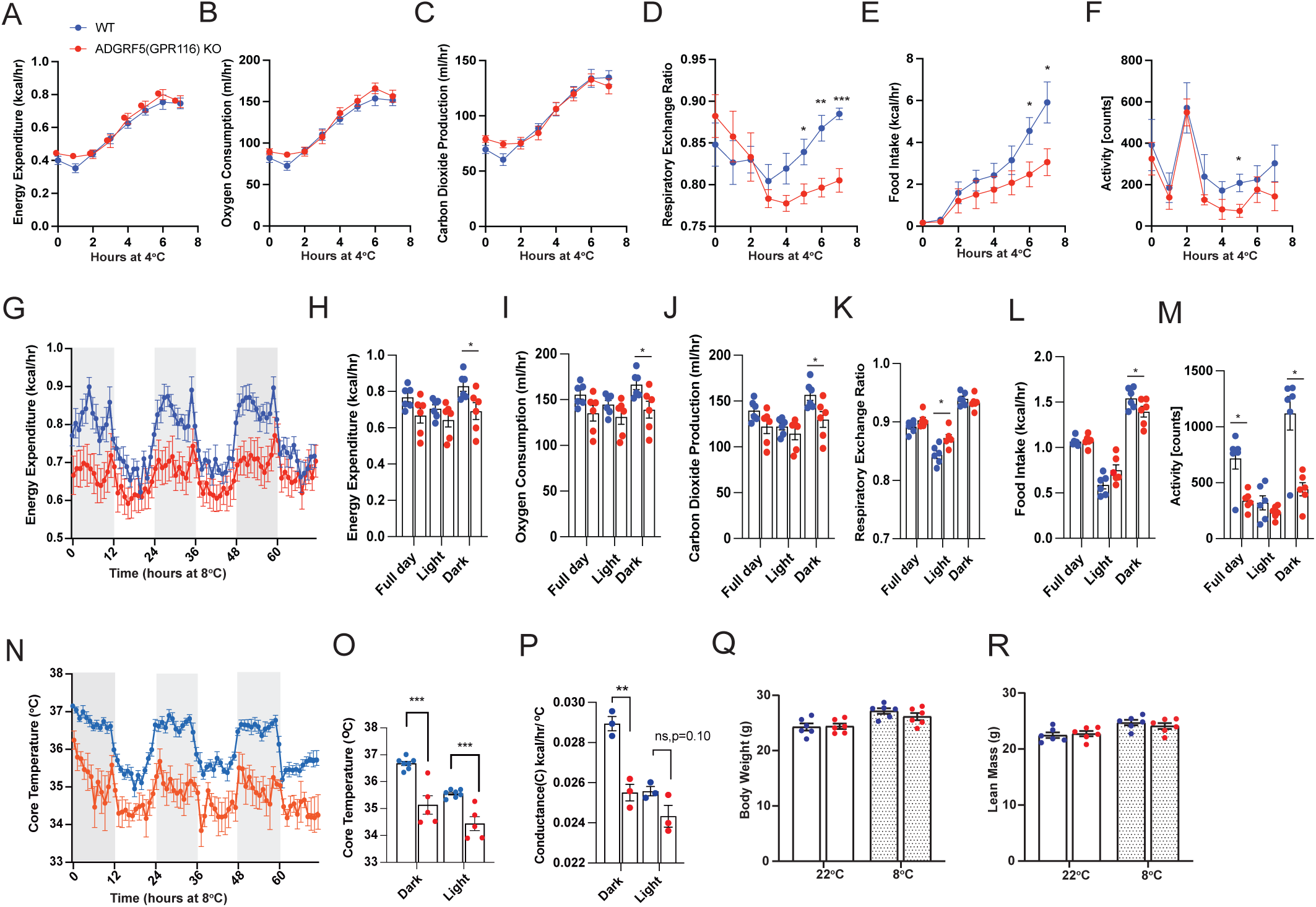
Global deletion of *Adgrf5(Gpr116*) impairs thermogenic energy expenditure during prolonged cold exposure. Indirect calorimetry measurement in ADGRF5(GPR116) KO mice and WT controls in metabolic cages at 4°C over 8 hours (A-F). Mice shown are males, C57BL6N, 9-10 weeks old, n=5-6 mice per group. (G-M) Indirect calorimetry measurement in ADGRF5(GPR116) KO mice and WT controls, in metabolic cages at 8°C for 14 days. The last 3 days are shown. Mice shown are males, C57BL6N, 11-12 weeks old, n=6 mice per group. (N-O) Core body temperature, (P) conductance, in ADGRF5(GPR116) KO mice and WT controls, exposed at 8°C for 14 days. The last 3 days are shown. (Q) Body weight, (R) Lean mass. All mice were single- caged. Statistical analysis was performed using CalR. Group differences were assessed using general linear modeling with or without ANCOVA, depending on body weight or lean mass differences, as implemented in the CalR platform. Data shown are means + SEM.* p-value <0.05, ** p-value< 0.01, *** p-value< 0.001. Grey shaded shapes indicate the Dark phase.

To examine whether ADGRF5(GPR116) KO may have experienced increased heat loss, as an underlying reason for lower core body T_m_ we calculated conduntance, which we found to be lower in the ADGRF5(GPR116) KO compared to the WT controls in the dark phase and similar between the genotypes in the light phase (Fig. 3O), suggesting that impaired heat production rather than increased heat loss contributed to the lower core body Tm in the ADGRF5(GPR116) KO. Since ADGRF5(GPR116) KO mice exchibited lower core body Tm in the dark phase, when activity levels where different between the genotypes, but also in the light phase when there was no observed differences in activity levels, we can expect that lower thermogenesis in the ADGRF5(GPR116) KO mice can be attributed to lower non-shivering and shivering thermogenesis, supporting an impaired BAT function in the ADGRF5(GPR116) KO mice. Finally, we observed no differences in body weight and lean mass between the genotypes (Fig. 3Q, 3R), supporting a non-mass effect in EE differences upon cold exposure.

### 3.4 Adipocyte Adgrf5(Gpr116) deletion does not account for impaired thermoregulation in ADGRF5(GPR116) KO mice

To directly assess the capacity for non-shivering thermogenesis (NST) after prolonged cold adaptation, we administered CL316,243 (CL)—a selective β3-adrenergic agonist—to mice housed at 8°C for 7 days. Both WT and ADGRF5(GPR116) KO mice exhibited an increase in core body temperature following CL stimulation (Fig. 4A), indicating preserved β3-adrenergic responsiveness in brown adipocytes. However, baseline core temperature remained lower in ADGRF5(GPR116) KO mice, and their thermogenic response to CL did not reach WT levels, suggesting a partial NST impairment during prolonged cold stress.

**Figure 4:**
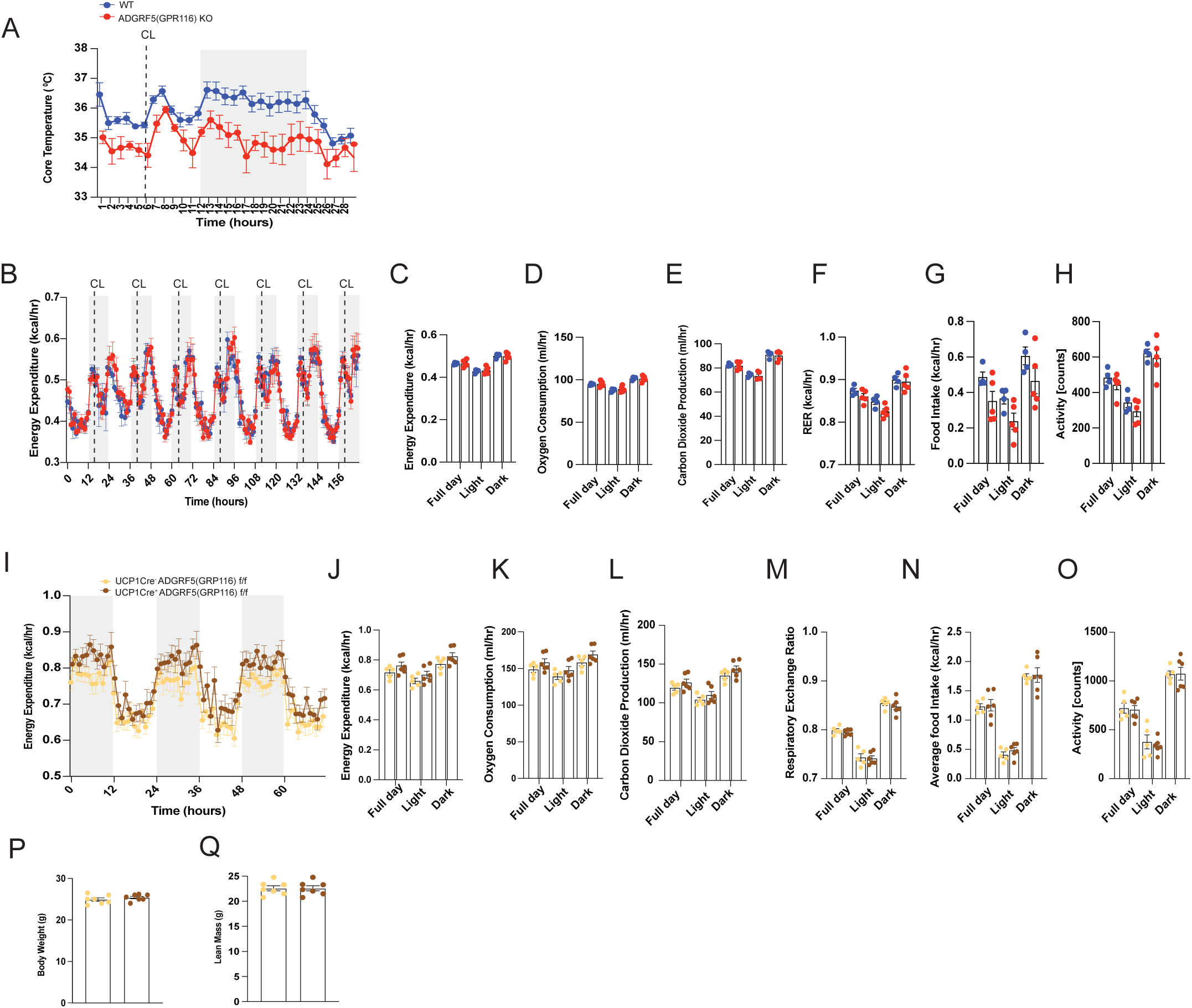
Adgrf5(Gpr116) deletion in brown adipocytes does not impair thermogenesis during β3-adrenergic activation or chronic cold adaptation. (A) Core body temperature of ADGRF5(GPR116) KO mice and WT controls kept for 14 days at 8°C. At day 12 mice were injected i.p with CL316,243 (CL). (B-H) Indirect calorimetry parameters were measured in ADGRF5(GPR116) KO mice and WT controls, which were injected i.p with CL in metabolic cages, once daily during the dark phase, for 7 consecutive days. (I-O) Indirect calorimetry parameters measured in UCP1Cre+ ADGRF5(GRP116) f/f mice and controls UCP1Cre- ADGRF5(GRP116) f/f mice, exposed at 8°C for 14 days. (P) Body weight and (Q) Lean mass of UCP1Cre- ADGRF5(GRP116) f/f and UCP1Cre+ ADGRF5(GRP116) f/f after 14 days at 8°C. Grey shaded areas on graphs indicate the Dark phase. Mice were single-caged. Mice were males, 11-12 weeks old. Statistical analysis was performed using CalR. Group differences were assessed using general linear modeling with or without ANCOVA, depending on body weight or lean mass differences, as implemented in the CalR platform. Statistical analysis was performed using CalR. Group differences were assessed using general linear modeling with or without ANCOVA, depending on body weight or lean mass differences, as implemented in the CalR platform. Data shown are means + SEM.

To further investigate brown adipocyte function in the absence of Adgrf5(Gpr116), we treated WT and ADGRF5(GPR116) KO mice with daily CL injections for 7 days under room temperature conditions. Both genotypes showed similar increases in EE, VO₂, VCO₂, and RER, with no significant differences in food intake or locomotor activity (Fig. 4B–4H). Additionally, brown adipocyte-specific deletion of ADGRF5(GPR116) using Ucp1-Cre (Ucp1-Cre+; ADGRF5(GPR116) f/f) did not impair thermogenic adaptation to prolonged cold exposure. These mice displayed comparable increases in EE, VO₂, VCO₂, RER, food intake, and activity to their Cre-negative littermate controls (Fig. 4I– 4O) and no difference in body weight and lean mass (Fig. 4P,4Q).

Together, these results indicate that the NST defect observed in cold-challenged ADGRF5(GPR116) KO mice is not attributable to loss of Adgrf5(Gpr116) in adipocytes. To rule out a neuroendocrine origin of the defect, we measured circulating levels of norepinephrine and thyroid hormone (T3)—two essential regulators of thermogenesis—and found no genotype-dependent differences (Suppl. Fig. 4A, 4B). Similarly, tyrosine hydroxylase (TH) expression in BAT, a marker of sympathetic innervation, was comparable between ADGRF5(GPR116) KO and WT mice (Suppl. Fig. 4C), indicating preserved neuronal input. Serum lipid profiles were also unchanged between genotypes (Suppl. Fig. 4D-4F), further supporting intact systemic responses to cold. Importantly, serum lactate levels were not elevated in ADGRF5(GPR116) KO mice compared to WT controls (Suppl. Fig. 4G), indicating normal systemic oxygenation and absence of hypoxia. This is consistent with published data showing that pulmonary dysfunction and emphysema in ADGRF5(GPR116) KO mice only manifest after 18 weeks of age.

Collectively, these data suggest that prolonged cold exposure in ADGRF5(GPR116) KO mice leads to maladaptive tissue remodeling that impairs the brown adipocyte microenvironment or paracrine signaling, ultimately limiting full thermogenic activation despite intact neuroendocrine and adipocyte-autonomous function.

### 3.5. Inducible deletion of Adgrf5(Gpr116) in endothelial cells impairs thermoregulation upon prolonged cold exposure without affecting angiogenesis

Since the thermogenic impairment in global ADGRF5(GPR116) KO mice could not be attributed to adipocyte-autonomous mechanisms or systemic neuroendocrine defects, we next investigated the role of *Adgrf5(Gpr116)* specifically in ECs—the primary site of its expression across multiple tissues, including BAT.

To this end, we generated an inducible endothelial-specific knockout model by crossing Cdh5-CreERT2 mice with ADGRF5(GPR116) f/f mice. This approach enabled deletion of *Adgrf5(Gpr116*) in adult ECs, thereby avoiding developmental compensation. Two weeks after tamoxifen administration, we confirmed efficient deletion of *Adgrf5(Gpr116*) from BAT ECs (Fig. 5A), and subsequently exposed the mice to 8°C for 7 days.

**Figure 5:**
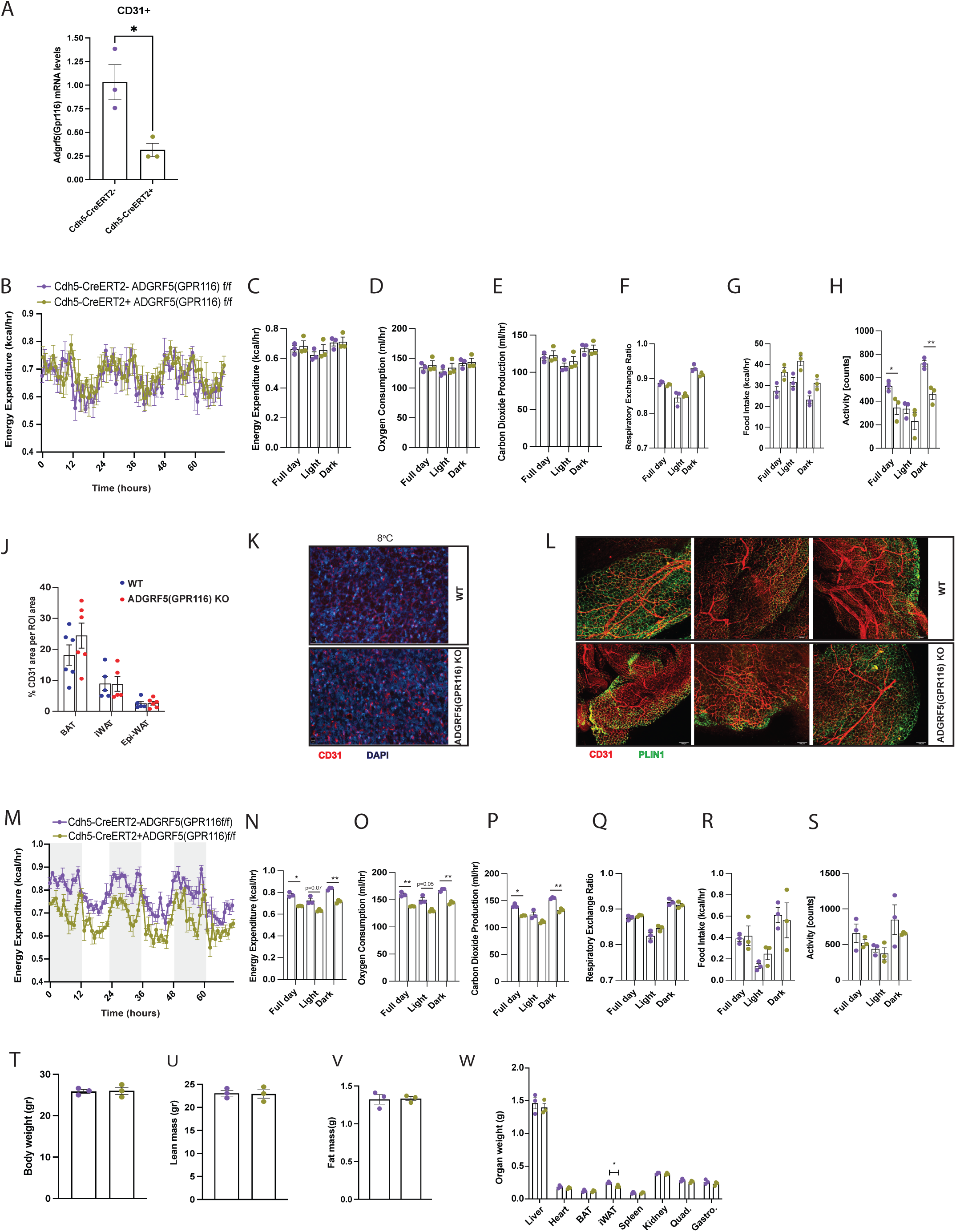
Inducible endothelial deletion of Adgrf5(Gpr116) impaired cold adaptation in response to prolonged cold without affecting angiogenesis. (A) qPCR quantification of *Adgrf5(Gpr116)* in BAT extracted ECs (CD31+) two weeks after tamoxifen injections in Cdh5CreERT2+ADGRF5(GPR116) f/f and Cdh5CreERT2- ADGRF5(GPR116) f/f mice. (B-H) Indirect calorimetry parameters of Cdh5CreERT2+ADGRF5(GPR116) f/f and Cdh5CreERT2-ADGRF5(GPR116) f/f mice housed 1 week at 8°C. Cold was initiated 2 weeks after the tamoxifen injections. (J) Immunofluorescent staining quantification of CD31 in 2D slides of indicated tissues from WT and global ADGRF5(GPR116) KO mice. (K) Representative immunofluorescent images for CD31 staining in BAT, at 8°C for 14 days, of indicated genotypes. (L) Immunofluorescent staining of CD31 and perilipin 1 (PLIN1) (3D staining) on inguinal WAT (iWAT) from WT and global ADGRF5(GPR116) KO mice. (M-S) Indirect calorimetry parameters of Cdh5CreERT+ADGRF5(GPR116) f/f and Cdh5CreERT-ADGRF5(GPR116) f/f mice were measured after 14 days, at 8°C, but they received tamoxifen injection after 1 week at 8°C. (T) Body weight, (U) Lean mass, (V) Fat mass, and (W) organ weights of Cdh5CreERT+ADGRF5(GPR116) f/f and Cdh5CreERT-ADGRF5(GPR116) f/f mice housed for 2 weeks at 8°C. Quad. is the quadriceps, gastro. is gastrocnemius. Mice were singly caged. Mice were females, 10- 12 weeks old. Statistical analysis was performed using CalR. Group differences were assessed using general linear modeling with or without ANCOVA, depending on body weight or lean mass differences, as implemented in the CalR platform. Data shown are means + SEM. .* p-value <0.05, ** p-value< 0.01, *** p-value< 0.001. Grey shaded shapes indicate the Dark phase.

During this cold challenge, knockout and control mice showed no differences in energy EE, VO₂, VCO₂, RER, or food intake (Fig. 5B–5G). However, physical activity was reduced in Cdh5-CreERT2+ ADGRF5(GPR116) f/f mice compared to controls (Fig. 5H). These data suggested that the deletion of endothelial Adgrf5(Gpr116) before cold exposure did not impair the acute thermogenic adaptation. Given that functional angiogenesis is critical from the onset of cold exposure to support thermogenesis, we hypothesized that Adgrf5(Gpr116) deletion did not compromise endothelial cell proliferation or new vessel formation. Moreover, this angiogenic remodeling may have partially restored Adgrf5(Gpr116) expression in newly formed endothelial cells, thereby compensating for the initial deletion. Consistent with this, BAT and WAT CD31 expression and the number of large vessels (quantified in iWAT, due to BAT’s dense microvascular network) were comparable between genotypes in the global ADGRF5(GPR116) KO (Fig. 5J-5L), indicating that endothelial Adgrf5(Gpr116) is not required for cold-induced angiogenesis. Nevertheless, the inducible deletion model may allow re-expression of *Adgrf5(Gpr116)* in newly generated endothelial cells during the cold challenge, potentially masking phenotypic effects during this early adaptive phase. To specifically evaluate the role of endothelial *Adgrf5(Gpr116*) in maintaining thermogenesis after the angiogenic phase, we injected tamoxifen into mice that had already undergone 7 days of cold exposure (8°C) —timing the gene deletion to the maintenance phase thermogenic response, as opposed to BAT expansion. In this context, endothelial-specific deletion of *Adgrf5(Gpr116*) led to a marked reduction in EE, VO₂, and VCO₂ (Fig. 5M–5P), without significant changes in RER or food intake (Fig. 5Q–5R), and still showing a trend toward decreased nighttime activity (Fig. 5S). Body composition was not different between the genotypes (Fig. 5T-5V).

Importantly, heart and spleen weights were comparable between genotypes (Fig.5W), suggesting that the increased heart and spleen mass observed in the global *Adgrf5 (Gpr116)* knockout is unlikely to account for the reduction in EE observed during cold exposure. While iWAT weight was significantly lower in Cdh5-CreERT2⁺ Adgrf5(Gpr116)f/f mice compared to controls (Fig. 5W), the difference was relatively modest, and total fat mass composition remained comparable between genotypes (Fig. 5V). This mild reduction in inguinal WAT (iWAT) weight points to a potential role for endothelial Adgrf5(Gpr116) in WAT remodeling, reminiscent of previously reported adipogenesis defects in Ap2–Cre–driven Adgrf5(Gpr116) deletion models (62). Since Ap2 is expressed in both adipocytes and adjacent endothelial cells, our data suggest that the previously observed adipogenic impairment may have a partial endothelial origin.

Together, these results demonstrate that endothelial Adgrf5(Gpr116) is dispensable for cold-induced angiogenesis, but plays a critical role in maintaining non-shivering thermogenesis during prolonged cold exposure.

### 3.6 Altered cell composition and intercellular communications in Adgrf5(Gpr116)-deficient BAT after prolonged cold exposure

The reduction in EE observed in both the constitutive global ADGRF5(GPR116) KO mice and the inducible endothelial-specific knockout mice (Cdh5-CreERT2+ ADGRF5(GPR116) f/f) upon prolonged cold exposure—despite normal physical activity and preserved angiogenesis—highlights a non-redundant, angiogenesis-independent role of Adgrf5(Gpr116) in thermogenic adaptation. Our prior experiments also ruled out adipocyte-autonomous and neuroendocrine causes, pointing toward a local microenvironmental mechanism.

To investigate alterations in BAT cell composition and intercellular communication, we performed single-nucleus RNA-seq analysis of BAT from mice with global deletion of Adgrf5 (Gpr116). Although these transcriptomic data were obtained from the constitutive global knockout model, the marked phenotypic similarity to the endothelial-specific knockout—particularly the cold-induced thermogenic defect— strongly suggests that the observed molecular changes primarily reflect endothelial dysfunction. We therefore leveraged this dataset to explore how loss of Adgrf5 (Gpr116) affects the transcriptional identity of key BAT cell populations and disrupts intercellular communication networks.

In both WT and ADGRF5(GPR116) KO BAT we sequenced a similar number of nuclei, suggesting an equal representation of both genotypes. In both genotypes and ambient temperature conditions, we identified the same cell populations, BAd, WAd, ECs (ArtECs, CapECs, LECs), MuralC (including VSCMs and pericytes), MyoC and MyogProg, Macro, SC, Prol.Prog-Fibro. (Fig. 6A). At both 22°C and 8°C ADGRF5(GPR116) KO BAT showed approx. 15 % less brown adipocytes compared to WT BAT, and it had 6% more WAd specifically in cold-exposed conditions, indicating that the differences in the numbers of pro-thermogenic (BAd) and anti-thermogenic (WAd) adipocytes favored a phenotype of impaired thermogenesis upon cold exposure. In addition, the ADGRF5(GPR116) KO mice showed a slight decrease in the number of APCs either at 22°C (-5%) or at 8°C (-1.4%) and a slight increase in the number of Prolf.Prog-Fibro. mainly at 8°C (+ 0.4%) overall, suggesting lower availability of adipogenic progenitors and a shift to profibrotic progenitors in BAT upon Adgrf5(Gpr116) deletion. We observed a striking increase of MyoProg in ADGRF5(GPR116) KO in both 22°C (+ 13%) and 8°C (+ 12%) and MyoC at 8°C (+ 11%), possibly suggesting a compensatory increase of myogenic lineage. The ADGRF5(GPR116) KO had 4% more ArtECs at 22°C and showed no further increase in the ArtECs numbers upon cold exposure, as opposed to the WT BAT, which showed a 4% more ArtECs after cold exposure. This could be indicative of an endothelial dysfunction, which becomes only evident upon cold stress. We did not observe any differences in the numbers of CapECs, LECs or MuralC between the genotypes (Fig. 6B).

**Figure 6:**
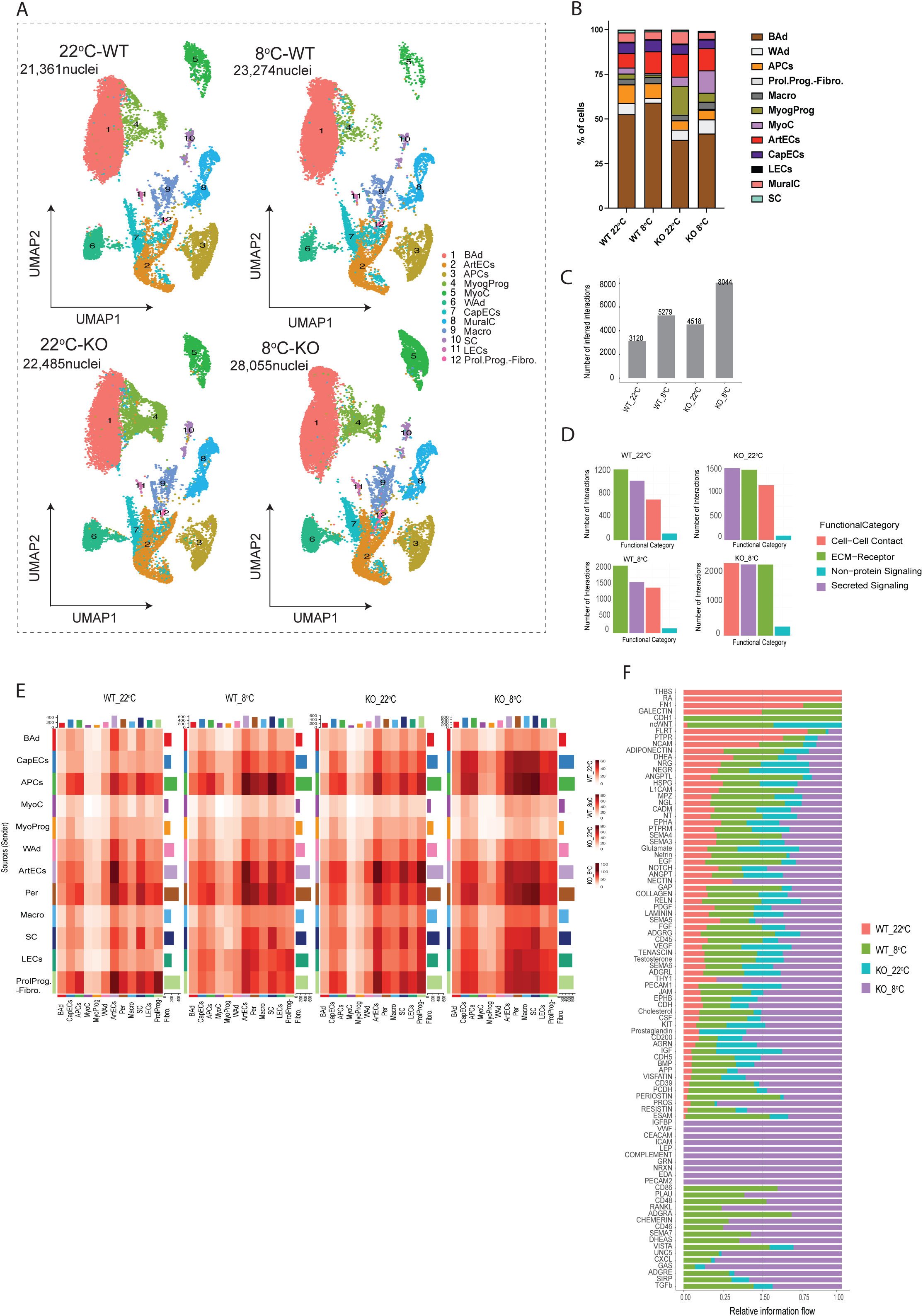
Effect of Adgrf5(Gpr116) deletion on BAT cellular composition and intercellular interactions upon cold exposure. (A) UMAP plots showing clustering of nuclei from WT and ADGRF5(GPR116) KO BAT at room temperature 22°C and prolonged cold exposure (8°C, 14 days). (B) Percentage of nuclei per cell type clusters, corresponding to (A). Barplots showing the number of cell-cell interactions (C) per condition, genotype, and (D) per functional group, per condition, genotype in BAT. (E) Heatmaps showing the interaction score between indicated cell types, as senders or receivers, at indicated conditions. (F) Bar plots showing the relative information flow for significant communication pathways between all BAT cell types, at the indicated conditions. KO is global ADGRF5(GPR116KO). Cell-cell interactions were calculated with the CellChat algorithm.

Pathway analysis using CellChat revealed that cold-exposed ADGRF5(GPR116) KO BAT exhibited a distinct remodeling of intercellular communication networks. We observed that the intercellular interactions were the highest in the KO BAT at 8°C compared to all other conditions (Fig. 6C). This was a result of increased interactions in all three functional categories (Cell-Cell contact, Secreted signaling, ECM-receptor and Non-protein Signaling) (Fig. 6D). Consistent with the crucial role of endothelial Adgrf5(Gpr116) in BAT tissue function during prolonged cold exposure the Adgrf5(Gpr116) deletion in BAT led to stronger interactions between ECs and stromovascular fraction (Fig. 6E). Specifically, KO-exclusive enrichment of VWF, ICAM, CEACAM, PECAM2, and IGFBP pathways suggests endothelial stress, altered adhesion, and possibly increased barrier permeability. Upregulation of pro-fibrotic and matrix-associated signals such as GRN and EDA indicated a dysfunctional ECM remodeling in ADGRF5(GPR116) KO BAT. In addition, we observed broader enrichment of pathways not specific to the KO but significantly upregulated, including NECTIN, CDH (cadherin), ESAM, and THY1—further supporting disrupted cell-cell junctions and compromised endothelial integrity. Inflammatory and immune-related cues such as PROSTAGLANDIN, COMPLEMENT, RESISTIN, CD46, CD200, and CXCL chemokines were also more prominent, reflecting increased inflammatory status in ADGRF5(GPR116) KO BAT. Moreover, alterations in signaling related to metabolic (LEP, Visfatin), neurovascular (NRXN, UNC5), and apoptotic or survival pathways (PLAU, RANKL, S1PR) pointed toward a complex microenvironmental imbalance (Fig. 6E). Collectively, these changes reinforce the notion that loss of Adgrf5(Gpr116) in ECs promotes maladaptive crosstalk and ECM remodeling, impairing the functional support of brown adipocytes during thermogenic stress.

### 3.7 BAT endothelial cells with Adgrf5(Gpr116) deletion show a transcriptional signature consistent with endothelial to mesenchymal transition and fibro-inflammation

To gain insight into the molecular pathways altered by Adgrf5(Gpr116) deletion in BAT endothelial cells during prolonged cold exposure, we performed Gene Set Enrichment Analysis (GSEA) on the single-nuclei transcriptomic data from ArtECs and CapECs subsets. LECs had a low number of nuclei, making it a small population to give meaningful results in the GSEA analysis (Extended File 4). We observed an ectopic upregulation of muscle contractility and cytoskletal remodeling specifically for KO cold ECs, alongside a downregulation of ATP synthesis and mitochondrial function, suggesting a metabolically stressed endothelial phenotype (Fig. 7A). This was confirmed by an ectopic transcriptional signature of sarcomeric and contractile proteins such as *Acta1, Des, Myh7, Myot*, actin binding and cytoskeletal remodeling, such as *Actn3, Actn,2 Synpo2, Lmod2, Shroom1-3* (Fig. 7B). Given that Adgrf5(Gpr116) is known to signal via Gαq and Rho GTPases (63, 64)—key regulators of actin cytoskeleton dynamics—we examined the expression of Rho GTPase signaling components in BAT ECs. Although not all genes were uniformly regulated, Adgrf5(Gpr116)-deficient ECs exhibited a distinct transcriptional signature affecting multiple nodes of the Rho–ROCK–actomyosin axis. This included differential expression of small GTPases (*RhoA, RhoB, RhoF, Rac1*), RhoGAPs and RhoGEFs (*Arhgap29, Dlc1, Arhgap17, Vav2, Dock7*), and downstream effectors involved in contractility (*Rock1, Myl9, Synpo2, Shroom1–3*). These transcriptional alterations suggest aberrant regulation of actin stress fibers and impaired integration of mechanical cues, a state associated with compromised endothelial identity and barrier integrity (65) (Fig. 7B). Indeed, tight junction markers such as *Tjp1 (ZO-1), Ocln, Cldn5, Esam* were reduced in KO ECs, whereas markers indicating increased permeability to leukocytes, such as *Sele, Cxcl12, Vcam1* were increased in KO ECs (Fig. 7B).

**Figure 7:**
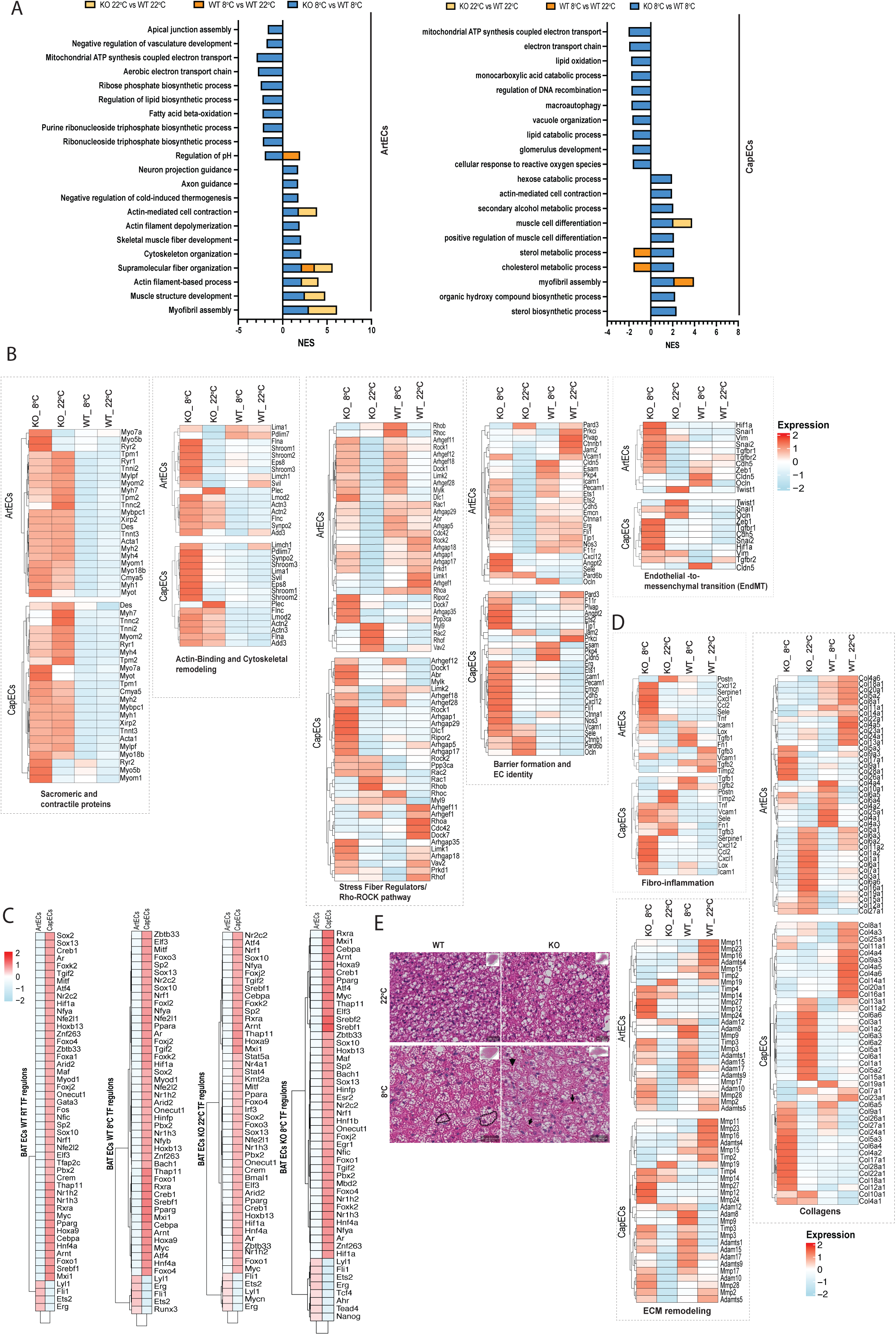
Transcriptional signature in ADGRF5(GPR116) KO BAT ECs supports EndMT activation and fibro-inflammation. (A) GSEA pathways analysis for the indicated conditions. NES scores of the top 10 significantly up- and down-regulated pathways are shown for the comparisons. (B, D) Heatmaps showing expression levels of indicated genes, organised in the indicated functional groups. (C) Heatmaps of regulons analysis for the indicated conditions. (E) H&E staining of BAT tissues for indicated conditions. Data are average expression per cluster shown produced with single-nuclei RNAseq from BAT of global deletion ADGRF5(GPR116KO) (here KO) and WT mice.

These features are hallmarks of early endothelial-to-mesenchymal transition (EndMT), prompting us to further examine this process. Indeed, KO ECs showed robust upregulation of canonical EndMT transcription factors (*Snai1, Snai2, Zeb1, Hif1a*), alongside mesenchymal markers (*Vim, Tagln, Tgfbr2*), indicating active transcriptional reprogramming toward a mesenchymal phenotype (Fig. 7B). Transcriptional regulon analysis confirmed this shift, revealing specific enrichment of EndMT-associated regulons under cold KO conditions, including Egr1, a known downstream effector of TGF-β signaling that promotes mesenchymal gene expression (66), and Tcf4 and Tead4, key mediators of Wnt/β-catenin and Hippo/YAP pathways respectively (67) — both central drivers of EndMT. In the KO RT conditions, the activation of Nr4a1 and Bmal1(Arntl) was particularly notable (Fig. 7C). Although Nr4a1 and Bmal1 are primarily circadian regulators, emerging evidence indicates they also modulate TGF-β/Smad signaling and vascular inflammation in fibrotic contexts (66, 68, 69). Their activation in cold-exposed Adgrf5(Gpr116)-deficient BAT ECs may thus reflect early engagement of profibrotic signaling, aligning with the transition toward an EndMT-like phenotype.

While EndMT has been described in WAT, particularly under obesogenic and inflammatory conditions such as high-fat diet, its role in BAT remains unexplored. In WAT, inflammatory cytokines (e.g., TGF-β, TNF-α, IL-1β), oxidative stress, and hypoxia promote endothelial reprogramming that contributes to fibrosis and adipose dysfunction (70). Given the EndMT-like features observed in Adgrf5(Gpr116)-deficient BAT ECs, we next assessed pro-fibrotic transcriptional phenotype and actively contributed to ECM remodeling. Consistent with this, collagen gene expression analysis revealed widespread induction of fibrotic ECM components in Adgrf5(Gpr116)-deficient ECs. Both ArtECs and CapECs showed robust upregulation of canonical fibrillar collagens (*Col1a1, Col1a2, Col3a1*), associated with interstitial fibrosis and matrix stiffening. Additional increases in *Col5a1–3* and *Col6a1–3* further support excessive ECM assembly and altered fibrillogenesis. Induction of remodeling- associated collagens (*Col14a1, Col16a1, Col22a1, Col24a1*) suggests broad architectural reorganization of the interstitial matrix. In contrast, expression of basement membrane–associated collagens (*Col4a1–a6*) was reduced or unchanged, indicating that barrier-specific ECM components are not maintained, while interstitial matrix is being aberrantly expanded (Fig. 7D). In parrallel, matrix-degrading and remodeling enzymes, including *Mmp12*, *Mmp27*, *Mmp14*, were induced in the KO BAT ECs, suggesting ongoing ECM turnover and tissue remodeling (Fig. 7D). This was accompanied by a fibro-inflammatory transcriptional profile marked by upregulation of cytokines (*Tnf*, *Ccl2*, *Cxcl1*, *Cxcl12*) and leukocyte adhesion molecules (*Sele*, *Icam1*, *Vcam1*), indicative of increased inflammatory cell recruitment capacity. Finally, elevated levels of *Tgfb1–3, Fn1, Serpine1*, and *Postn*—especially in capillary ECs— further support the engagement of profibrotic signaling pathways and active matrix reorganization (Fig. 7D).

These transcriptomic changes were mirrored histologically by the appearance of distorted brown adipocyte structures and dilated vessels in cold-exposed Adgrf5(Gpr116) KO BAT (Fig. 7E), indicating that endothelial Adgrf5(Gpr116) deletion disrupts both vascular integrity and the adipocyte microenvironment. Together, these findings suggest that Adgrf5(Gpr116)-deficient BAT ECs initiate a fibro-inflammatory program characterized by excessive ECM deposition, barrier destabilization, and inflammatory activation, ultimately impairing vascular–adipocyte crosstalk and thermogenic tissue remodeling.

## 4. Discussion

Our study reveals a previously unrecognized role for the aGPCR Adgrf5(Gpr116) in preserving endothelial identity and suppressing fibro-inflammatory remodeling in BAT. We show that Adgrf5(Gpr116) deletion in BAT ECs triggers a fibro-inflammatory transcriptional program consistent with endothelial-to-mesenchymal transition (EndMT), characterized by cytoskeletal remodeling, induction of stress fiber components, upregulation of collagens and matrix metalloproteinases, and loss of endothelial identity and barrier genes. These changes occur in the absence of angiogenesis defects, highlighting a novel, angiogenesis-independent role for Adgrf5(Gpr116) in preserving endothelial identity and restraining maladaptive matrix remodeling. Our findings establish that BAT endothelial cells are not passive conduits but active contributors to tissue fibrosis, capable of modulating thermogenic output through paracrine extracellular matrix remodeling. This places Adgrf5(Gpr116) at the center of a newly appreciated endothelial mechanism that regulates the balance between functional thermogenesis and pathological remodeling in metabolically active tissues.

ECs are highly sensitive to mechanical forces such as shear stress, stretch, and pressure, and respond through specialized mechanosensory systems that regulate cytoskeletal dynamics, gene expression, and barrier function(71). In brown adipose tissue (BAT), the vasculature consists primarily of a dense capillary network with relatively few large arteries or arterioles. During cold exposure, BAT perfusion increases significantly to meet the high metabolic demand, which is expected to elevate local shear stress within the capillary bed. While increased perfusion has been linked to angiogenic remodeling in cold-acclimated BAT, the broader impact of mechanical stimuli on endothelial function—and how this influences thermogenic tissue remodeling—remains incompletely understood. Our finding that Adgrf5(Gpr116) deletion causes endothelial dysfunction only during prolonged cold exposure, but not under basal or acute cold conditions, strongly supports a role for Adgrf5(Gpr116) in enabling endothelial adaptation to sustained perfusion and shear stress. Importantly, the absence of angiogenesis defects in our model suggests that Adgrf5(Gpr116) is dispensable for vessel formation but is crucial for the stabilization and functional maintenance of the remodeled vasculature. This endothelial adaptation, in turn, appears to influence adipocyte function through paracrine matrix remodeling, highlighting a previously unrecognized link between vascular mechanosensing and thermogenic regulation.

Endothelial-to-mesenchymal transition (EndMT) is a context-specific cellular program by which ECs acquire mesenchymal, contractile, and matrix-remodeling characteristics. It is broadly categorized into three types: Type I, which occurs transiently during development; Type II, associated with fibrosis and tissue injury; and Type III, observed in tumor progression (72). In our model, the loss of Adgrf5(Gpr116) in BAT ECs leads to a transcriptional program characteristic of Type II EndMT, including the upregulation of stress fiber components, actomyosin regulators, cytoskeletal remodeling genes (*Shroom1–3*, *Synpo2*, *Myl9*), and classical EndMT transcription factors (*Snai1*, *Snai2*, *Zeb1*, *Hif1a*), as well as the loss of junctional and endothelial identity genes (*Cdh5*, *Tjp1*, *Cldn5*). This EndMT phenotype depends on cold exposure, suggesting that mechanical or metabolic stress triggers endothelial reprogramming in the absence of Adgrf5(Gpr116)-mediated regulation. The convergence of phenotypes in both constitutive and inducible models underscores the role of Adgrf5(Gpr116) in restraining pathological stress-induced Type II EndMT in adult BAT tissue. Interestingly, Adgrl4(ELTD1), another aGPCR highly expressed in BAT endothelium, has been shown to work together with Adgrf5(Gpr116) (73) and to induce an endothelial-to-mesenchymal transition (EndMT) toward a myofibroblast phenotype (74), suggesting that functionally related aGPCRs may co-regulate endothelial identity and plasticity in thermogenic fat.

Tissue expansion inherently stresses the endothelium: in BAT, cold drives capillary growth and perfusion, while in WAT, obesity enlarges adipose depots. Chronic obesity has been linked to accelerated EndMT in WAT prior to fibrosis (70), suggesting that EndMT may facilitate ECM restructuring during physiological expansion. However, when combined with persistent inflammatory cues—as seen in insulin resistance or diabetes—EndMT shifts into a fibro-inflammatory phenotype, producing collagen and matrix remodeling enzymes (70, 75). Importantly, inducible endothelial-specific deletion of Adgrf5(Gpr116) displayed reduced WAT pad size under prolonged cold exposure, implicating endothelial Adgrf5(Gpr116) in normal adipose expansion—a finding paralleling earlier observations that adipose-specific (aP2-Cre) (62) but not adiponectin-Cre deletion(76) of Adgrf5(Gpr116) impairs fat pad expansion and insulin sensitivity. Because aP2-Cre is also highly expressed in ECs, next to adipocytes and adiponectin-Cre does not recapitulate the lipodystrophic phenotype, these results underscore an endothelial-specific role for Adgrf5(Gpr116) in adipogenesis, likely through regulating a reversible, remodeling-focused EndMT. Thus, Adgrf5(Gpr116) appears to function as a molecular gatekeeper, preventing EndMT from progressing into maladaptive fibrosis once tissue expansion demands have been met.

Fibrotic remodeling of brown adipose tissue (BAT) is increasingly recognized as a barrier to thermogenic plasticity, especially in aging, obesity, and metabolic disease. Recent work has highlighted that fibro-inflammation, characterized by concurrent extracellular matrix (ECM) accumulation and low-grade inflammation, impairs BAT function and is associated with reduced thermogenic capacity during diet-induced obesity (21). While adipocytes, stromal cells, and immune cells have been proposed as sources of fibrotic signals, the role of the endothelium in driving fibro-inflammatory remodeling in BAT has been largely overlooked. Our data demonstrate that loss of Adgrf5(Gpr116) in BAT ECs leads to robust upregulation of ECM and remodeling genes, including *Col1a1*, *Col4a1*, *Mmp12*, and *Mmp27*, under cold exposure, independent of angiogenesis defects. These changes point to a non-angiogenic, endothelial-driven program of fibro-inflammation that disrupts adipose homeostasis. Importantly, the fibrotic signature in Adgrf5(Gpr116)-deficient BAT emerged only during prolonged cold exposure, suggesting that fibro-inflammation in BAT is a dynamic and stress-inducible process, not simply a byproduct of aging or obesity. This is consistent with emerging models in which mechanical or metabolic stress can initiate EndMT-like transitions in endothelial cells, leading to maladaptive ECM remodeling. Thus, Adgrf5(Gpr116)-deficient ECs are not passive bystanders but active drivers of fibro-inflammatory remodeling, linking impaired endothelial plasticity to the suppression of thermogenic competence in cold-adapted BAT.

Adgrf5(Gpr116) has been previously implicated in maintaining endothelial integrity in barrier-rich tissues, including the lung and the brain(59). Studies of global or endothelial-specific Adgrf5(Gpr116) knockout mice have reported vascular leakage, pulmonary surfactant imbalance, and age-dependent emphysema—phenotypes that typically become apparent in mice older than 18 weeks (36, 59). Similarly, deletion of other endothelial-enriched receptors such as Adgrl4 (ELTD1) has been linked to blood–brain barrier (BBB) dysfunction, hemorrhage, or vascular malformations, but these often require either developmental deletion or severe inflammatory or hypoxic stimuli to manifest (59, 73). In contrast, our cold-exposure studies were performed in mice aged 9–12 weeks, and we did not observe any lung pathology, BBB breakdown, or signs of hemorrhage, even in global knockout animals. This suggests that cold exposure triggers a unique, context-specific vulnerability in BAT endothelium that is not shared by other vascular beds under similar conditions. Notably, the endothelial-specific inducible knockout of Adgrf5(Gpr116) phenocopies the global constitutive knockout with regard to defective thermogenic maintenance during prolonged cold exposure, despite some divergence in broader systemic phenotypes. For example, increased spleen and heart weights were observed only in the global knockout, not in the inducible endothelial-specific model. This reinforces the conclusion that the core BAT thermogenic phenotype arises primarily from endothelial loss of Adgrf5(Gpr116), and is not secondary to general systemic dysfunction or developmental abnormalities. Taken together, these findings highlight that Adgrf(Gpr116) function is highly tissue- and stimulus-specific, playing a non-redundant role in maintaining BAT endothelial identity and matrix homeostasis specifically in response to chronic thermogenic stress, rather than under steady-state conditions or in other vascular territories.

Together, our work suggests that endothelial Adgrf5(Gpr116) acts as an important modulator linking mechanosensation to vascular stability and thermogenic tissue functional integrity, offering new insight into the complex interplay between endothelium and adipose tissue homeostasis.

## Acknowledgements

We would like to thank the Timo Müller lab and the Siegfried Ussar lab, specifically Andreas Israel, for access to facilities, equipment, and material. We thank Kerstin Lohr for all the support with animal protocols and animal licenses. We thank Do Khanh Linh Nguyen, MSc student in our lab, for assistance with sample measurements. We thank Jiangyan Yu, from the University of Würzburg, for her advice on integration methods and data analysis. We thank Inti De La Rosa Velazquez and Ugur Dura, from the Helmholtz Munich Core Facility Genomics (CF-GEN), for support with single-nuclei RNA sequencing and data analysis. We thank Annette Feuchtinger, Judith Bushe, and Richard Linder, from the Helmholtz Munich Core Facility Pathology and Tissue Analytics (CF-PTA) for their services on tissue staining and pathological evaluation.

## Authors’ Contributions

RE.M, V.K, designed and performed experiments, analyzed data, and prepared the manuscript figures. R.K, L.S, S.G. MY.J, S.Hi, M.H, E,K, AK.J, AT.K, provided material and performed experiments and provided input which improved the duirng the manuscript preparation. K.D, F.M provided conceptual input and experimental advice, J.H, M.K, A.P, J.H, D.W ,S.H provided straregic input, critically reviewed the manuscript, supervised students and participated in meetings, where the project progress and concepts development were discussed. A.G conceptually conceived the study, supervised the project development, analyzed the data, and wrote the manuscript. All authors read and approved the manuscript.

---

**Supplementary Figure 1:**
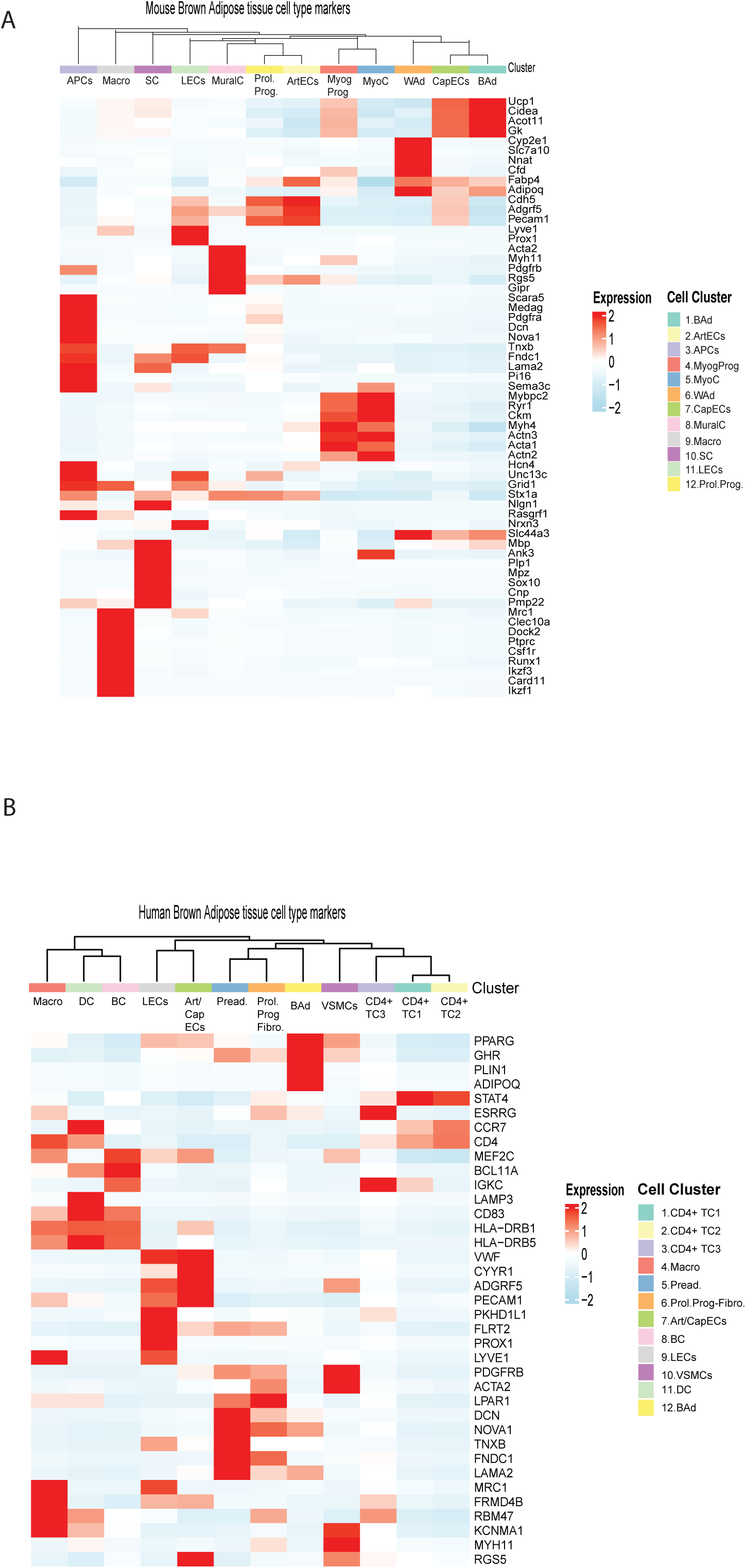
Canonical markers of cell clusters in (A) mouse and (B) human BAT. Mouse BAT is our dataset generated in this study. Human BAT is from the published study Sun W. et al (47)

**Supplementary Figure 2:**
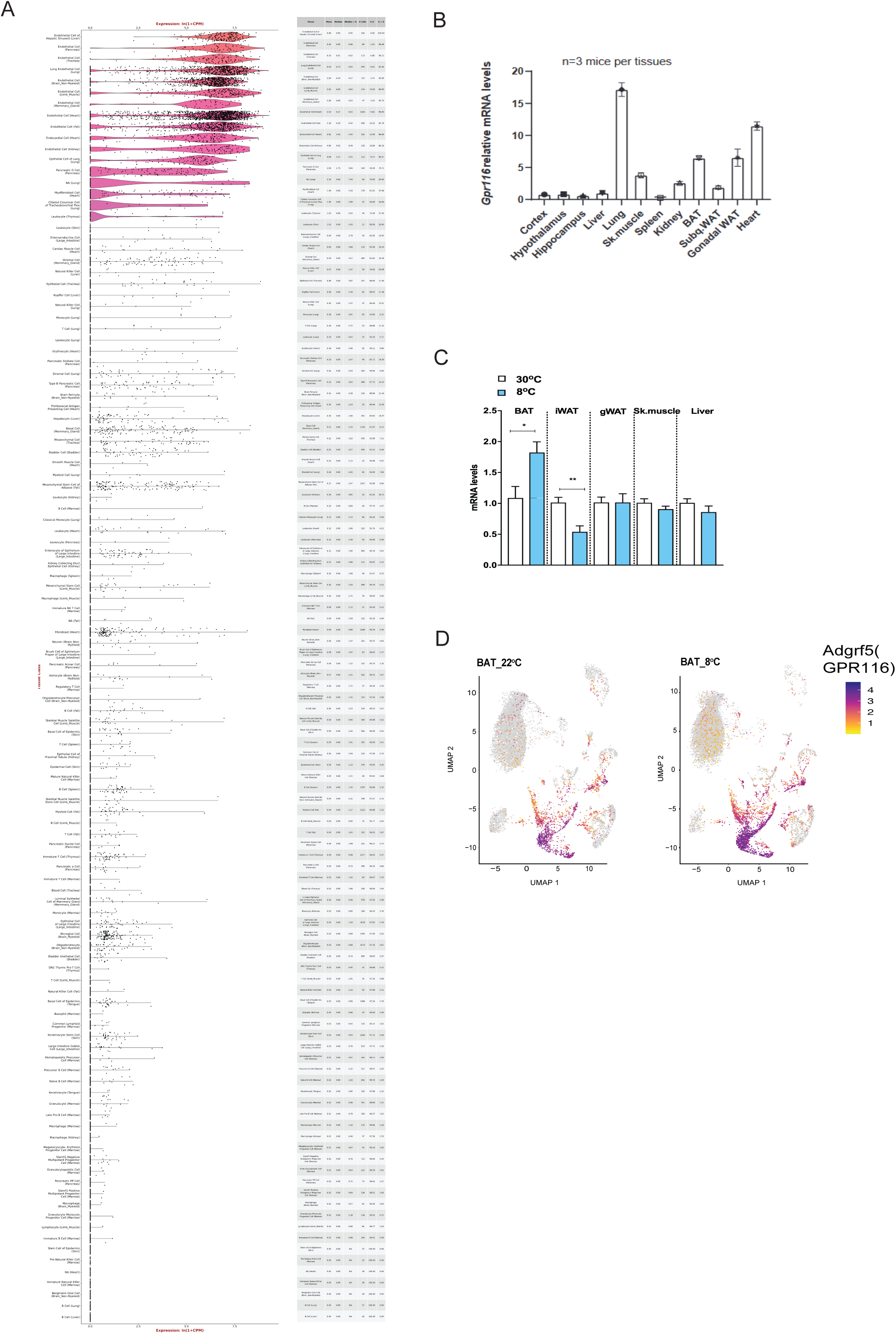
Cellular expression of Adgrf5(Gpr116) across all mouse tissues and regulation in BAT upon cold exposure. (A) Single-cell expression levels of Adgrf5(Gpr116) across different tissues. Image generated from the Tabula Muris project browser. (B) qPCR quantification of *Adgrf5(Gpr116)* at indicated tissue. Mice are males, C57Bl6N, 9 weeks old, n=3 per tissue. (C) qPCR quantification of *Adgrf5(Gpr116)* at the indicated tissues and ambient temperatures. Mice were acclimated at 30°C for 3 weeks and moved from 30°C for 8°C (after a gradual drop in ambient temperature) for 7 days. BAT is brown adipose tissue, iWAT is inguinal white adipose tissue, gWAT is gonadal white adipose tissue, Sk. Muscle is skeletal muscle and it is the gastrocnemius. Mice are males, C57Bl6N, 9 weeks old, n=3-4 per tissue. (D) Single-nucleus expression of *Adgrf5(Gpr116*) at indicated ambient temperatures in WT mice (C57Bl6N). Data generated from the single-nuclei RNAseq presented in this paper. Mice are males, C57Bl6N, 10-12 weeks old, n=3-4 per tissue. Data shown are means + SEM.* p-value <0.05, ** p-value< 0.01, *** p-value< 0.001. Statistics are Student’s t-test.

**Supplementary Figure 3:**
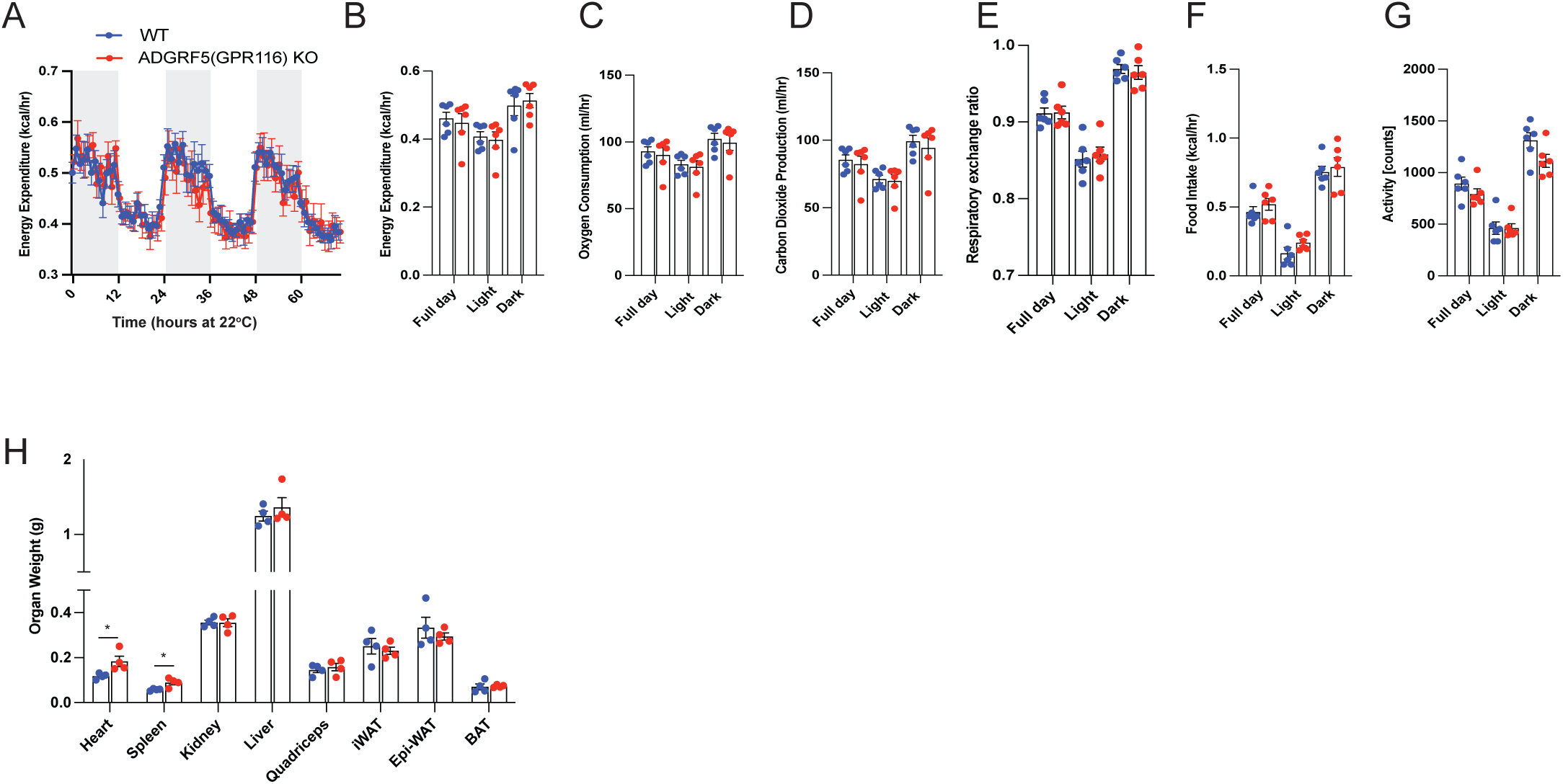
Characterization of metabolic phenotype of ADGRF5(GPR116) KO at room temperature (22°C). Indirect calorimetry measurement in ADGRF5(GPR116) KO mice and WT controls in metabolic cages at 22°C for 14 days. The last three days are shown (A-G). Mice are males, C57Bl6N, 10-12 weeks old, n=6. (H) Organ weights of the above ADGRF5(GPR116) KO mice and WT controls at the end of the study. n=4 per tissue. Data shown are means + SEM.* p-value <0.05, ** p-value< 0.01, *** p-value< 0.001. Statistics are Student’s t-test.

**Supplementary Figure 4:**
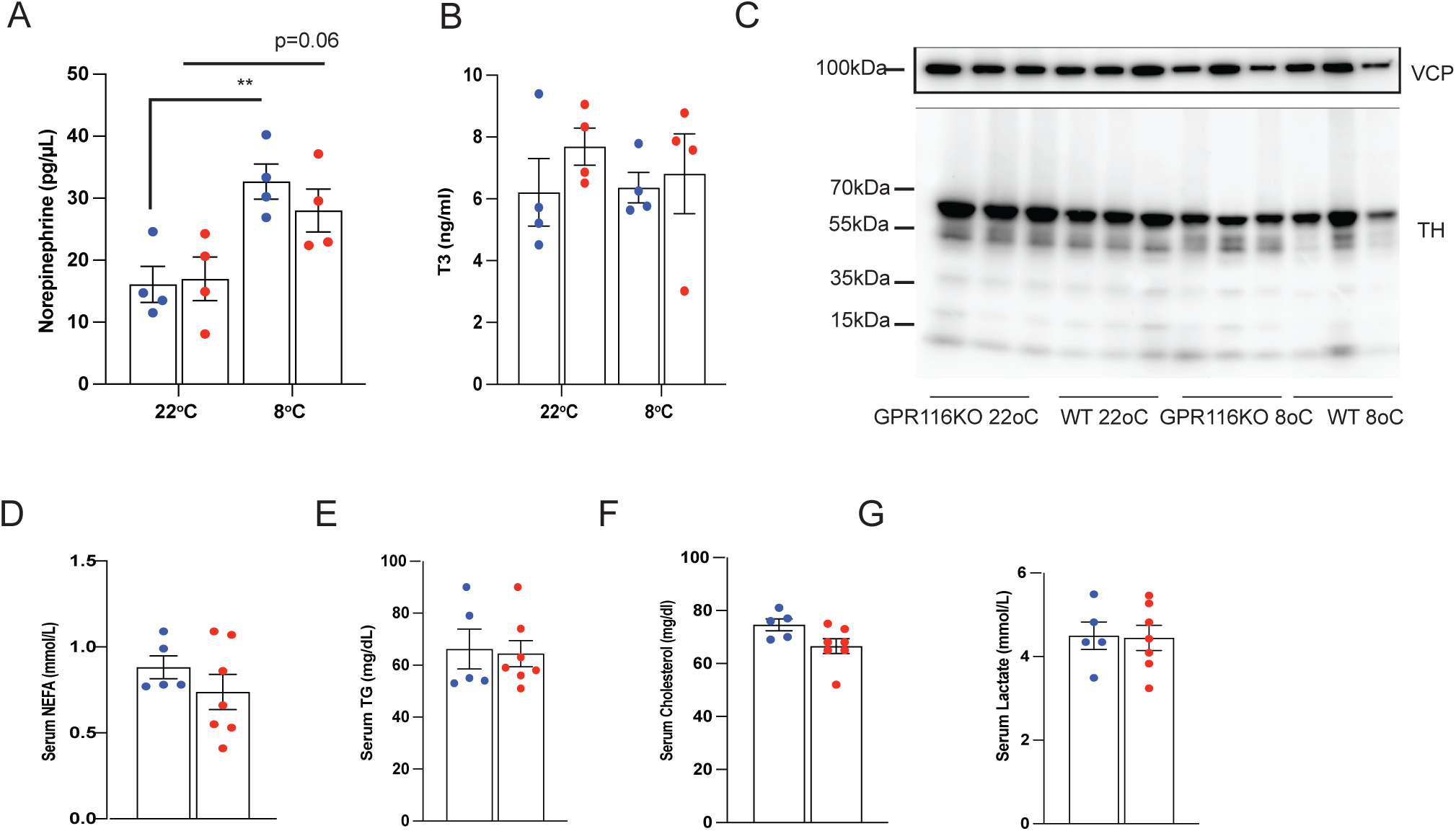
Neuroendocrine and systemic circulation parameters in ADGRF5(GPR116) KO mice. (A) HPLC quantification of plasma norepinephrine. (B) ELISA quantification of serum thyroid hormone (T3). (C) Western blot analysis of tyrosine hydroxylase (TH) and Vinculin (VCP) in BAT total cell lysates. (D-G) Serum analyser quantification of indicated serum parameters. Mice are males, C57Bl6N, 10- 12 weeks old, exposed to cold 8°C for 14 days. Data shown are means + SEM ** p- value< 0.01. Statistics are Student’s t-test. **Declaration of generative AI and AI-assisted technologies in the writing process** During the preparation of this work, the authors used OpenAI’s ChatGPT (2024 version) for troubleshooting and optimizing code during the analysis of single-nucleus transcriptomic data, as well as for improving clarity, grammar, and structure throughout the text. After using this tool, the authors reviewed and edited the content as needed and take full responsibility for the content of the publication.

**Supplementary Table 1.**
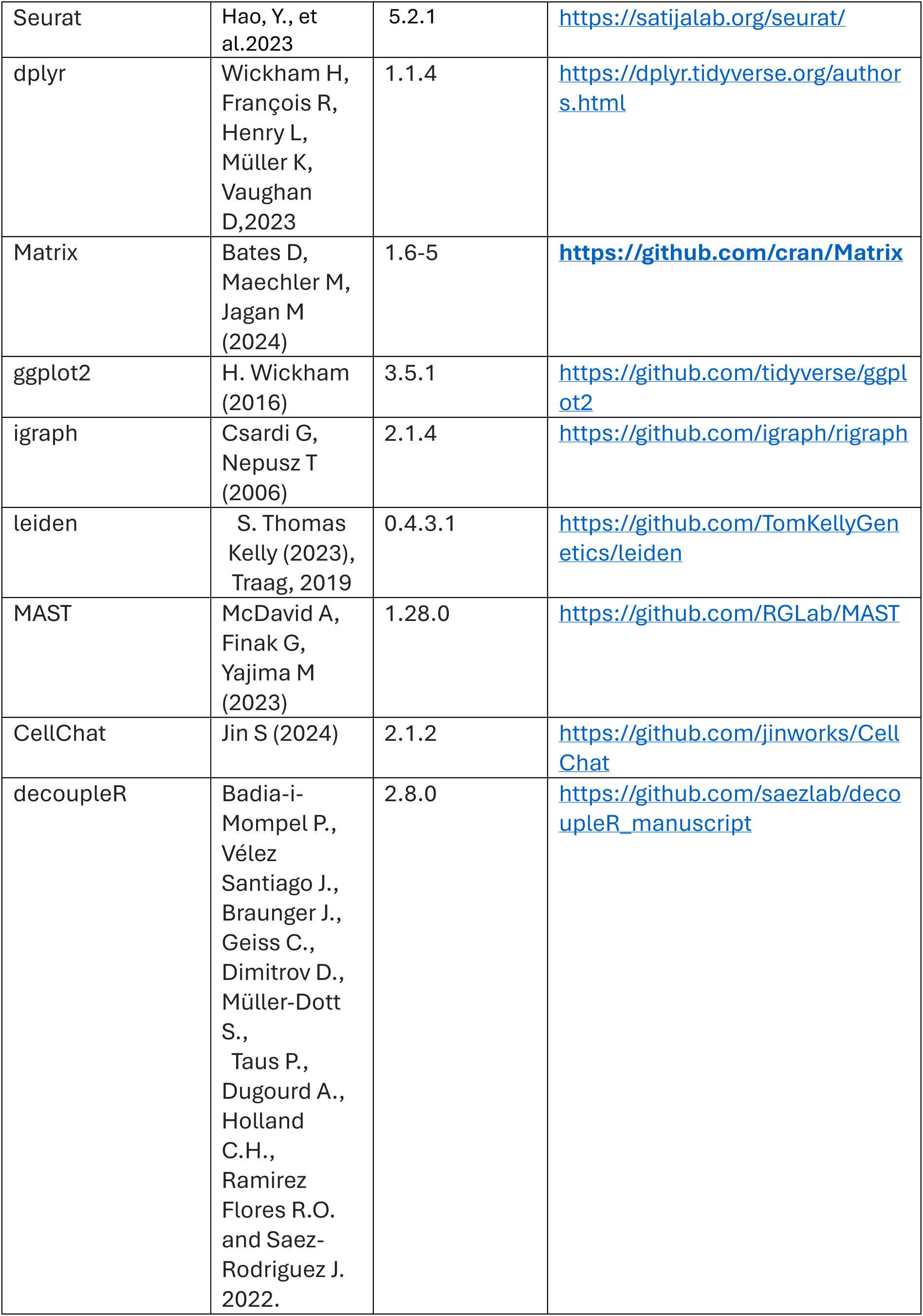
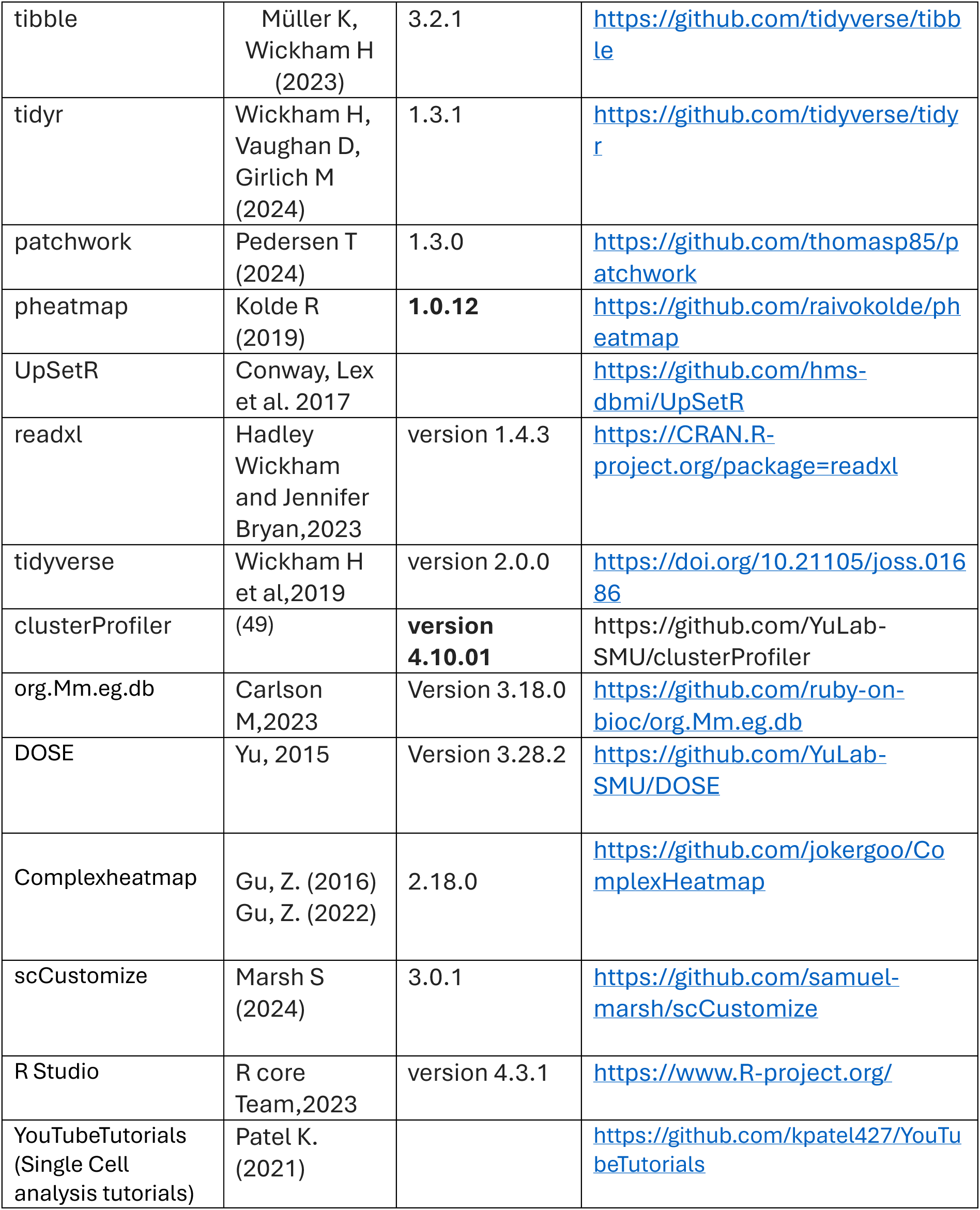
Software and algorithms.

